# PULSAR: a Foundation Model for Multi-scale and Multicellular Biology

**DOI:** 10.1101/2025.11.24.685470

**Authors:** Kuan Pang, Yanay Rosen, Kasia Kedzierska, Ziyuan He, Abhejit Rajagopal, Claire E. Gustafson, Grace Huynh, Jure Leskovec

**Author notes:** These authors contributed equally (Kuan Pang, Yanay Rosen).

## Abstract

Biology emerges from interactions across physical scales, where molecular interactions drive cellular states, which in turn orchestrate multicellular tissue functions that collectively define health and disease. However, current computational models are often constrained to single scales in isolation, failing to integrate the biology that emerges from lower to higher levels [1]. Here we present PULSAR (**P**atient **U**nderstanding **L**everaging **S**ingle-cell univers**A**l **R**epresentation), a multi-scale and multicellular foundation model architecture that explicitly enables information flow from genes to cells to multicellular systems. Applied to the human peripheral immune system, PULSAR extracts a unified donor representation that supports rapid disease classification, biomarker prediction, and forecasting of future clinical events, such as Rheumatoid arthritis onset. As a generative model, PULSAR enables the simulation of cytokine perturbation response across physical resolutions, while its interpretability reveals the key cell types driving disease. Overall, PULSAR opens new avenues for precision medicine by enabling computational reasoning that connects molecular biology to clinical phenotypes.

## 1 Main

Measuring, mapping, and modeling of multicellular systems has long been a goal of both experimental and computational biology [2–9]. Groups of billions of specialized cells in the human body coordinate to maintain homeostasis, respond to environmental challenges, and execute complex physiological programs that collectively define health and disease [10–12]. Developing an AI model that can quantify and simulate these multicellular systems is essential for understanding and predicting human physiology and pathology [1, 13, 14].

Accurately modeling multicellular biology requires reasoning across physical scales. Biology emerges from increasingly complex interactions: perturbations at the molecular level drive functional shifts in individual cells, which, in turn, alter cell-type composition and coordination to reshape system-level biology [5, 15]. Consequently, a robust understanding of an individual-level phenotype from a multicellular system cannot exist in isolation; it must be rooted in the lower-level molecular and cellular states that drive it. To achieve this understanding, we need to build digital models that integrate information from genes to cells to multicellular systems and capture emergent properties at the individual level.

Despite recent breakthroughs in generative AI for modeling molecular [16–20] and cellular data [21–25], achieving multi-scale integration remains elusive due to the uneven availability of data across physical scales, where data availability scales inversely with physical complexity. While raw molecular [26–28] and cellular profiles [29–31] are now abundant, high-quality datasets containing sets of related, interacting cells, sampled from a multicellular system, such as a tissue or donor, remain comparatively scarce [13, 32]. Consequently, naive approaches that attempt to model organismal phenotypes directly, without explicit grounding in lower-level mechanisms, suffer from a critical loss of inductive bias and a narrow view of biological variation. As a result, they require prohibitive amounts of training data to rediscover biological rules and ultimately yield only a shallow understanding of the system. Conversely, single-scale models restricted to a cel-lular or molecular view lack the systemic context needed to capture the complex interactions that emerge at the multicellular level. Overcoming this impasse demands a hierarchical architecture that learns progressively across scales, exploiting abundant unlabeled data and leveraging robust representations at the molecular and cellular levels to construct a flexible framework for multicellular reasoning.

Here we present PULSAR (**P**atient **U**nderstanding **L**everaging **S**ingle-cell univers**A**l **R**epresentation), a new multi-scale, multicellular foundation model architecture for integrating molecular, cellular, and multicellular understanding through a connected representation learning framework across physical layers. PULSAR uses specialized encoding models to learn representations at each physical scale while remaining flexible with respect to data modality, and connects the next physical scale using the learned embeddings from previous scales to inject a hierarchical inductive bias. The model integrates biological information at different resolutions: the molecular layer captures genomic properties, regulations, and interactions; the cell layer encodes cellular functional phenotypes; and multicellular aggregation preserves sample context and cellular composition.

We applied PULSAR to the human peripheral immune system, a uniquely compelling domain for modeling multicellular coordination. Peripheral blood mononuclear cells (PBMCs) provide a dynamic window into systemic health, reflecting both tissue-specific pathologies [33, 34] and whole-body physiological programs [35–37]. While clinically accessible, PBMCs exhibit significant variation in cell-type composition and transcriptomic states across age, sex, and disease, making them ideal biomarkers for precision medicine [38–41]. Supported by recent expansions in massive PBMC single-cell RNA sequencing (scRNA-seq) atlases [39, 42–44], this domain offers the population-scale data needed to train a foundation model of multicellular organization. scRNA-seq also provides an ideal modality for multi-scale modeling, as it simultaneously resolves transcript levels and cell type heterogeneity within a single sample.

We trained PULSAR on billions of transcript readouts from 36.2 million cells across 6,807 donors to extract multicellular information into a unified donor embedding. We demonstrate that this representation space captures donor-level phenotypes comparable across diverse studies and disease conditions, powering a vector database, `DONORxEMBED`, which enables reference mapping against 2,804 donors spanning inflammation landscapes. PULSAR’s zero-shot donor embedding accurately decodes plasma protein levels and can predict future clinical events, including Rheumatoid Arthritis conversion and flu vaccine responsiveness. Beyond prediction, PULSAR is generative: by manipulating the donor representation and decoding it to finer resolutions, we can simulate cytokine perturbations across physical scales. Crucially, the model remains interpretable, identifying the key cell types and gene programs driving disease progression. Together, PULSAR establishes a scalable framework for bridging multi-resolution biological readouts to donor-level phenotyping, opening new avenues for precision medicine and translational discovery.

## 2 Results

### 2.1 Modeling Multi-scale, Multicellular Biology with PULSAR

We design PULSAR to model multicellular systems through a hierarchy of three nested representation spaces, molecular, cellular, and multicellular, mirroring the natural organization of biological systems (Figure 1a). This multi-scale architecture enables information to flow progressively: molecular features inform cellular representations, which are then aggregated to capture multicellular organization, ultimately generating a unified donor-level embedding. Crucially, this design is conceptually agnostic to the specific foundation models employed at each scale.

**Figure 1.**
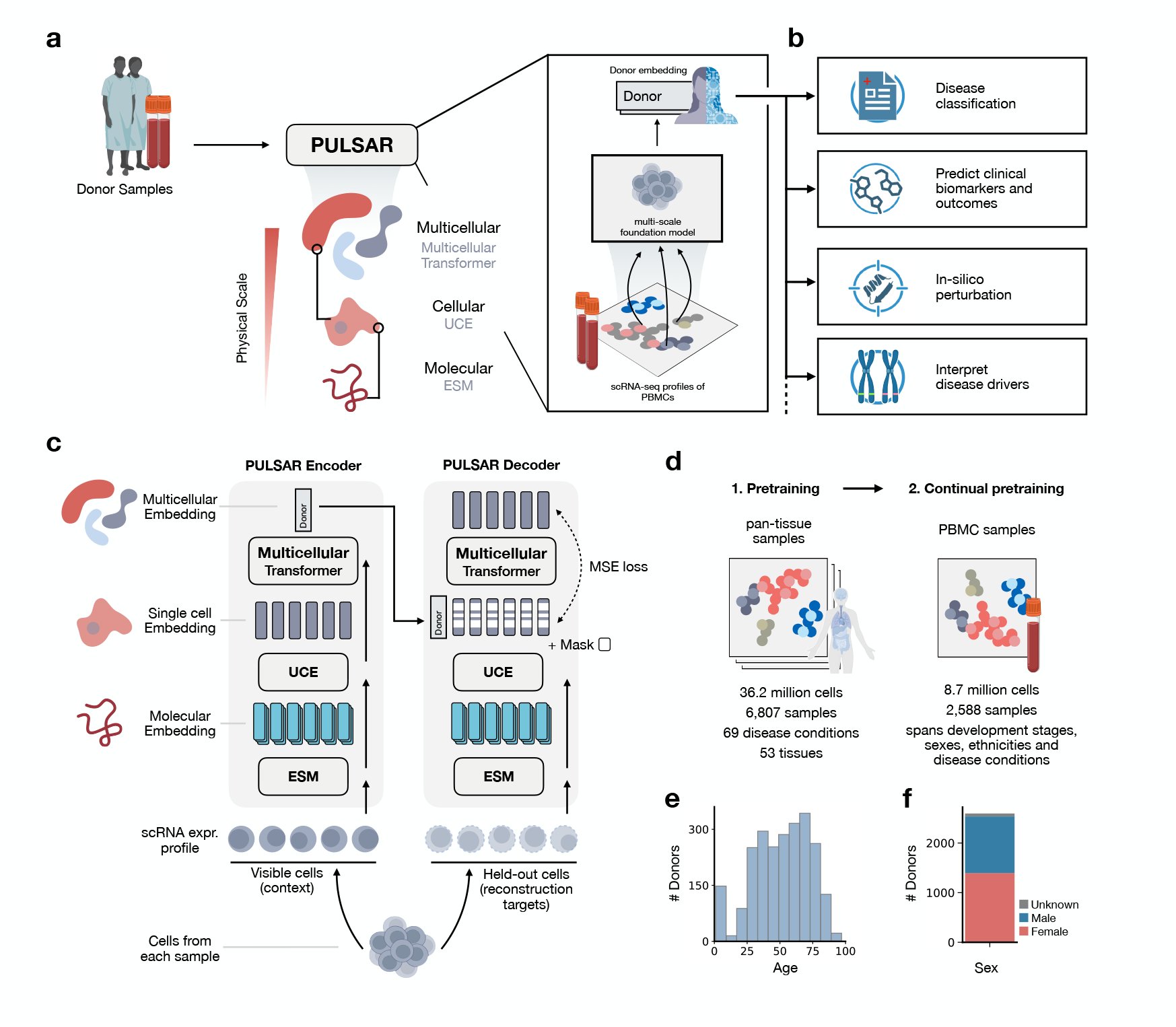
PULSAR connects physical scales of biology to learn donor-level representations. **(a)** Architecture of PULSAR, a multi-scale, multi-cellular foundation model that transforms a blood sample’s scRNA-seq profiles into a donor-level embedding. PULSAR progressively integrates molecular, cellular, and multicellular information, preserving the natural biological hierarchy. **(b)** Applications enabled by PULSAR, including donor disease classification, clinical biomarkers and outcomes prediction, in-silico perturbation simulation, and interpretation of disease drivers. **(c)** PULSAR’s self-supervised encoder-decoder architecture. The PULSAR Encoder uses ESM2 [19] for molecular embeddings, UCE [94] for single-cell representations leveraging molecular embeddings, and a Multicellular Transformer to generate donor embeddings by aggregating cell embeddings. The PULSAR Decoder reconstructs held-out cells using another Multicellular Transformer, and is trained with masking and Mean Squared Error (MSE) loss. **(d)** Two-phase pretraining strategy: initial pan-tissue pretraining on 36.2 million cells from 6,807 samples across 69 disease conditions and 53 tissues, followed by continual pretraining on 8.7 million cells from 2,588 PBMC samples spanning developmental stages, sexes, ethnicities, and disease conditions. **(e**,**f)** Training corpus demographics showing comprehensive age distribution (e) and balanced sex representation (f).

scRNA-seq is an ideal modality for multi-scale modeling, measuring gene expression profiles at single-cell resolution within a multicellular context. To train PULSAR using scRNA-seq data from circulating immune cells, we use three foundation models that encode the information at each scale (Figure 1b). At the molecular level, we leverage ESM2 [19], a protein language model trained on amino-acid sequences, to encode the functional properties of each gene’s protein product. At the cellular scale, PULSAR employs Universal Cell Embeddings (UCE) [21] to transform raw gene expression profiles into single-cell representations. UCE integrates the ESM2 representations with gene expression abundance, thereby capturing cell states that are robust to technical batch effects. Finally, at the multicellular scale, we introduce Multicellular Transformer, a new machine learning model that learns how collections of cells orchestrate to produce system-level phenotypes. Through self-attention [45], Multicellular Transformer contextualizes each donor’s immune profile by modeling both the composition and coordination of their cell populations while maintaining single-cell resolution. As a result, the donor embedding captures cell-specific alterations and compositional information about cell types.

PULSAR employs self-supervised learning to discover patterns in immune system organization (Methods 4.1). By relying solely on the intrinsic structure of biological data rather than human annotation, the model remains free of biases, allowing it to internalize native population-scale variations and learn biological rules that are strictly emergent from the data. To do this, we introduce a pretraining objective, Masked Cell Modeling, designed specifically for variable-sized, unordered sets of cells. The training architecture comprises two complementary networks (Figure 1c): PULSAR Encoder and PULSAR Decoder. For each donor, PULSAR first samples a subset of cells and passes them through the PULSAR Encoder to produce a single donor-level embedding. A separate subset of cells is then withheld and partially masked in their cell-embedding space: large fractions of their embedding dimensions are replaced with learnable masks. The PULSAR Decoder receives only the donor embedding and these masked cell embeddings, and is trained to reconstruct the missing features (Methods 4.2.2). By masking substantial portions of the cell embeddings, we force the model to learn robust cross-cellular dependencies rather than simple interpolation. This conditional reconstruction task also ensures that donor representations capture both compositional and functional properties of the sample.

We implement a two-phase training strategy to learn increasingly specialized representations (Figure 1d). Phase one (Pretraining) leverages 36.2 million cells from 6,807 donors across CELLxGENE [29, 46], encompassing diverse tissues and conditions. This pan-tissue pretraining phase establishes a broad understanding of cellular relationships and trains the model to encode sets of cells into embeddings. Phase two (Continual Pretraining) focuses on 8.7 million peripheral blood cells from 2,588 donors, carefully curated to represent all developmental stages and balanced demographics (Figure 1e,f). This stage refines the broadly learned representations toward peripheral blood–specific structure, improving sensitivity to rare immune cell subsets, and inter-individual variation critical for tasks.

### 2.2 PULSAR Enables Accurate Disease Classification Through Large-Scale Donor Reference Mapping

Inflammation-related diseases present significant diagnostic challenges due to heterogeneous clinical presentations and the absence of definitive biomarkers [47]. To evaluate PULSAR’s capacity in capturing disease-relevant donor-level signatures, we develop a reference mapping approach that leverages existing single-cell atlases for disease classification (Figure 2a). We compile an extensive reference cohort comprising 2,804 donors across 41 independent studies (Extended Data Table 1), totaling over 10 million PBMCs with paired diagnostic annotations (Figure 2b). We categorize these diseases into six classes that reflect their inflammatory landscape [48], encompassing a range of immune dysfunction and inflammatory states, including autoimmune diseases, acute inflammation, chronic inflammation, infections, malignancies (cancer), and healthy controls.

**Figure 2.**
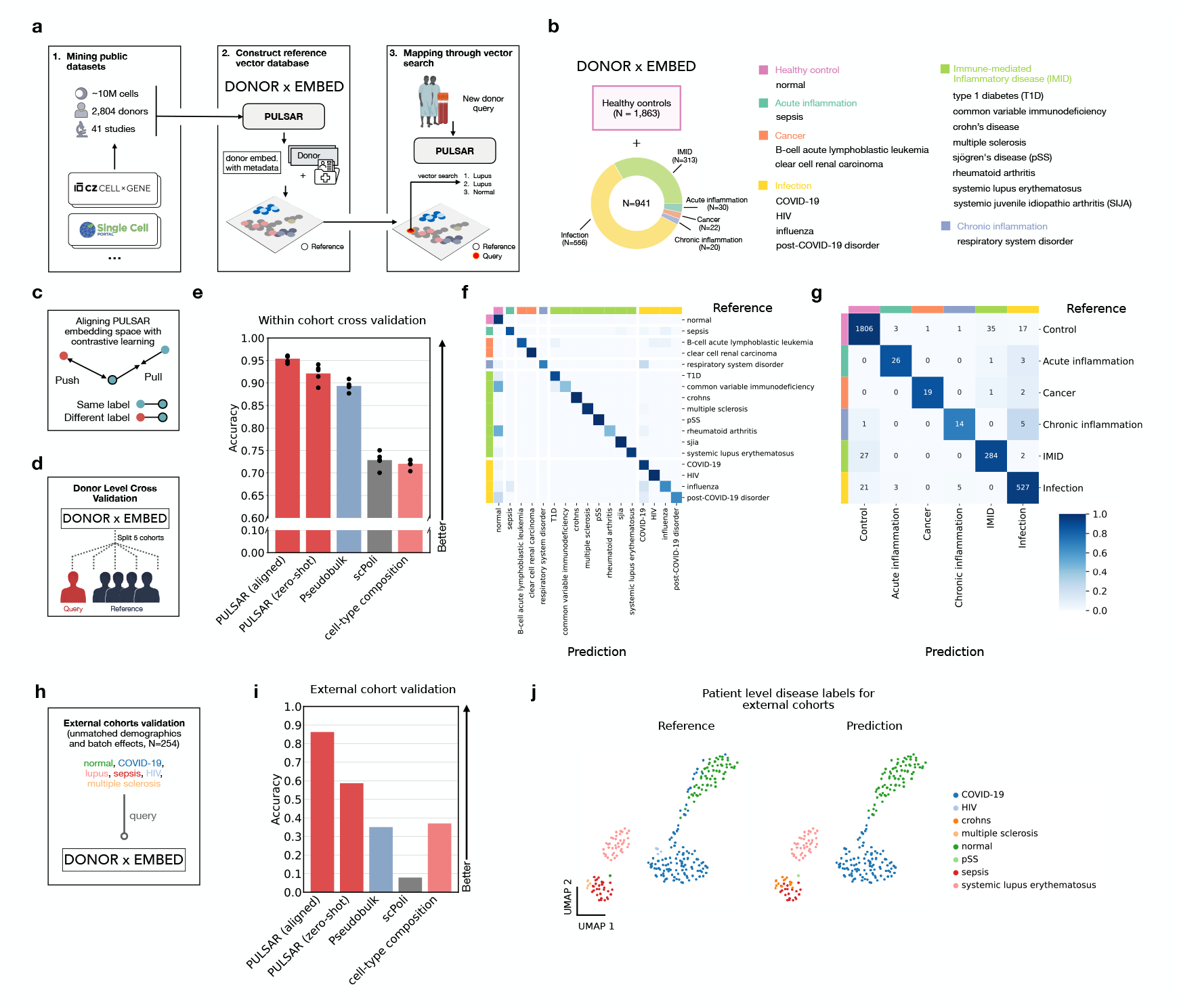
PULSAR enables accurate disease classification through large-scale donor reference mapping. **(a)** Workflow for disease classification using PULSAR. Public PBMC datasets containing paired single-cell RNA-seq and clinical data are processed to construct a reference vector database (`DONOR×EMBED`), enabling disease classification via *k*-nearest neighbors in the PULSAR embedding space. **(b)** Composition of the `DONOR×EMBED`reference database comprising 2,804 donors across 41 studies with over 10 million cells. Disease categories span immune-mediated inflammatory diseases (IMIDs), infections, malignancies (cancer), acute and chronic inflammations. **(c)** Contrastive alignment of the PULSAR donor embedding space. Donor embeddings produced by PULSAR are passed through a projection network and optimized with a contrastive loss that pulls together embeddings from donors with the same disease label (positive pairs) and pushes apart embeddings from donors with different diseases (negative pairs). **(d)** Donor-level cross-validation scheme ensuring complete separation of individuals between reference and query sets. **(e)** Classification performance comparison showing accuracy for aligned PULSAR, zero-shot PULSAR, Pesudo Bulk, scPoli [49], and cell-type composition baselines. Points represent each fold in cross-validation (*n* = 5 runs). **(f)** Full confusion matrix showing classification performance across all disease categories in the reference database. Color intensity represents the proportion of predictions for each true-predicted disease pair. **(g)** Aggregated confusion matrix for major immune-related disease categories. **(h)** External validation setup using independent cohorts with distinct patient populations, sequencing platforms, and batch effects (*n* = 254 donors). **(i)** External validation performance showing accuracy, demonstrating robust generalization of aligned PULSAR compared to baselines. **(j)** UMAP visualization of donor embeddings from external cohorts colored by predicted and reference disease labels.

To enable flexible, scalable disease classification, we constructed a searchable vector database, `DONOR×EMBED`, containing PULSAR-generated donor embeddings paired with clinical disease labels. This approach enables classification via *k*-nearest neighbors (kNN) search in the embedding space, allowing continuous reference expansion without re-training classifiers. To optimize the embedding space for disease discrimination, we also apply contrastive learning alignment to PULSAR using disease labels from each fold’s training set of 2243 samples (Figure 2c).

We evaluate performance using rigorous donor-level cross-validation within the cohort (Figure 2d). PULSAR is benchmarked against several baselines: the unaligned zero-shot embedding (PULSAR (zero-shot)), a gene expression profile baseline without single-cell resolution (Pseudobulk), a sample embedding baseline (scPoli [49]), and a cell-type composition (CTC) baseline, as many diseases show altered cell-type compositions [48, 50]. PULSAR with contrastive alignment achieved a state-of-the-art mean accuracy of 0.954 across disease categories, outperforming unaligned PULSAR embeddings (0.921), pseudobulk (0.893), scPoli embeddings (0.729), and CTC (0.720) (Figure 2e). The performance gap was particularly pronounced for macro F1 scores, where aligned PULSAR achieved 0.828 compared to 0.084 for CTC and 0.677 for pseudobulk (Supplementary Figure S1), indicating robust performance across imbalanced disease categories.

Detailed examination of the full confusion matrix reveals biologically meaningful classification patterns. PULSAR accurately discriminates between closely related conditions, for instance, distinguishing Systemic Lupus Erythematosus from Rheumatoid Arthritis despite shared autoimmune features (Figure 2f). The aggregated confusion matrix for major disease categories demonstrates near-perfect separation between healthy controls and disease states (Figure 2g), with primary misclassifications occurring within viral infections, where acute viral illnesses elicit a shared interferon-driven host programme that is largely pathogen-agnostic in peripheral blood [51, 52].

To assess generalization capability, we further evaluate PULSAR on completely independent external cohorts comprising distinct patient populations, sequencing assays, and clinical settings (Figure 2h). Without any fine-tuning on this external cohort, aligned PULSAR maintained accuracy of 0.862, compared to 0.370 for CTC approaches (Figure 2i). Macro F1 scores showed similar robustness, with PULSAR achieving 0.450 versus 0.094 for CTC methods (Supplementary Figure S2). This performance retention across technical and biological batch effects demonstrated that PULSAR learns fundamental disease signatures rather than dataset-specific patterns.

Visualization of the embedding space for external cohorts revealed high consistency between Reference and Prediction labels (Figure 2j). Further coherent disease clustering emerged without supervision of disease mechanisms. Infectious diseases (COVID-19, HIV) form distinct clusters reflecting pathogen-specific immune responses. Immune-mediated conditions (multiple sclerosis, Crohn’s disease, systemic lupus erythematosus) showed partial overlap consistent with shared inflammatory pathways, while maintaining disease-specific boundaries.

### 2.3 PULSAR Accurately Predicts Plasma Protein Levels

Proteins serve as direct molecular effectors of diverse biological processes, capturing dynamic pathophysiological changes and functioning as well-established clinical biomarkers [53]. To evaluate PULSAR’s zero-shot capacity to capture systemic physiological states, we develop a framework for predicting plasma proteomics from donor embeddings, in which PULSAR is neither trained on PBMC data from these donors nor on paired plasma proteomics.

We assemble a dataset of 436 blood samples collected from 176 donors with paired high-dimensional plasma proteomics (454 proteins) and PBMC scRNA-seq profiles [54], enabling direct assessment of transcriptome-proteome relationships (Figure 3a). The plasma proteomics panel comprises five panels covering major physiological domains: cardiovascular (*n* = 92), inflammation (*n* = 89), metabolism (*n* = 91), immune response (*n* = 92), and organ damage (*n* = 90). We employ donor-level cross-validation to ensure robust data split (Figure 3b), with PULSAR generating zero-shot donor representations that serve as input features for linear prediction models without any finetuning (Figure 3c, Methods 4.3.2).

**Figure 3.**
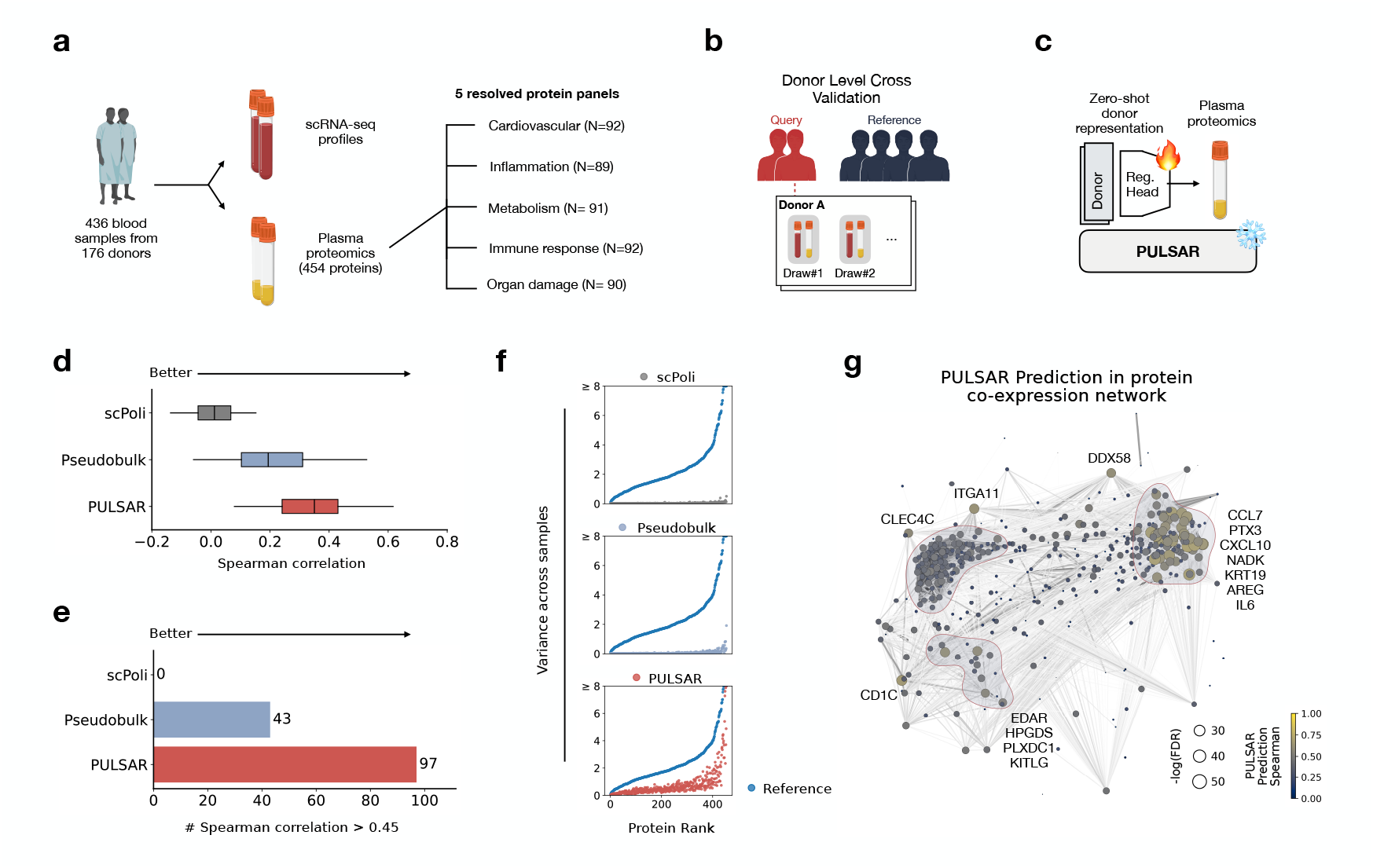
PULSAR enables accurate prediction of plasma proteomics from peripheral blood transcriptomes. **(a)** Schematic of the experimental approach. 436 blood samples from 176 donors were profiled using both scRNA-seq and plasma proteomics (454 proteins). Distribution of plasma proteins across five physiological domains: cardiovascular (*n* = 82), inflammation (*n* = 89), metabolism (*n* = 91), immune response (*n* = 92), and organ damage (*n* = 90). **(b)** Donor-level cross-validation scheme ensuring complete separation of individuals between reference and query sets. **(c)** Zero-shot prediction framework. Donor scRNA-seq profiles are processed through PULSAR to generate embeddings, which serve as features for plasma protein prediction models using regularized linear regression. **(d)** Spearman correlation between predicted and measured plasma protein levels across three embedding methods (scPoli [49], pseudobulk, PULSAR). Each box shows the interquartile range (IQR; 25^th^–75^th^ percentiles), the center line marks the median, and whiskers span the 5^th^–95^th^ percentiles; outliers beyond the whiskers are not shown. **(e)** Number of proteins achieving Spearman correlation ≥ 0.45. **(f)** Variance across donors is shown for each plasma protein, with proteins ranked along the *x*-axis by their empirical variance. Each panel overlays the variance predicted by one model (dots; scPoli, pseudobulk, or PULSAR, respectively, from top to bottom) on the same ordered set of proteins. **(g)** PULSAR prediction accuracy for individual proteins in protein co-expression network. Each point represents one protein, with size indicating −log_10_(FDR) and color showing prediction accuracy. Key immune and inflammatory markers are labeled.

Direct comparison of Spearman correlations (Spearman *r*) between predicted and measured protein levels revealed substantial improvements, with PULSAR achieving mean correlations of 0.339 for all proteins while pseudobulk (0.209) and scPoli (0.011) embeddings showed limited predictive capacity (Figure 3d, protein-level results shown in Supplementary Figure S5,S6,S7,S8,S9).

Quantitative assessment demonstrated that PULSAR achieved moderate correlations (Spearman *r* ≥ 0.45) for 97 proteins, compared to 43 for pseudobulk approaches and close to zero for scPoli (Figure 3e). This performance improvement was also consistent across all five protein panels, with Inflammation and Immune Response yielding higher performance improvement than other panels (Supplementary Figure S10). Further, a rank–variance analysis across samples (Figure 3f) showed that PULSAR yielded a smooth, intermediate-variance spectrum across the proteome, preserving a biologically plausible dynamic range.

To contextualize PULSAR’s prediction performance within the broader plasma proteome landscape, we visualize prediction correlation across an inferred protein co-expression network (Figure 3g, Methods 4.3.2). PULSAR demonstrates strong predictive performance across diverse functional clusters, including inflammatory mediators (IL6, PTX3, CXCL10) [55, 56], immune cell markers (CLEC4C, CD1C) [57, 58], and tissue damage indicators (KRT19, AREG) [59–61]. Notably, proteins within the same functional modules often show similar prediction correlation, suggesting that PULSAR captures coordinated biological programs. Some proteins with lower prediction accuracy tend to localize to network peripheries, potentially reflecting less interaction with circulating cells.

To identify proteins that benefit most from PULSAR’s single-cell–resolution representations, we jointly examine absolute prediction performance (PULSAR Spearman *r*) and improvement over pseudobulk (Δ(PULSAR, Pseudobulk)) (Supplementary Figure S11). Proteins above a performance threshold (PULSAR Spearman *r* ≥ 0.45) that also show large gains over pseudobulk (Δ ≥ 0.35) include markers of discrete immune cell subsets and signaling receptors (ZBTB16, LAT2, INPPL1, FKBP1B) [62–65] as well as mitochondrial and metabolic enzymes and DNA repair machinery (AIFM1, DECR1, GLO1, MGMT) [66–68]. Together, these targets illustrate how PULSAR leverages multicellular, cell-state–specific expression signatures that bulk-resolution profiles cannot capture.

### 2.4 PULSAR Predicts Rheumatoid Arthritis Onset and Flu Vaccine Response

We next investigate whether an individual’s baseline (steady-state) immune state is predictive of subsequent biological outcomes with zero-shot PULSAR embeddings. Prior studies have shown targeted immune readouts can forecast outcomes, such as response to flu vaccines [69, 70], but these approaches are task-specific and lack a single-cell-resolved context. We hypothesize that PULSAR embeddings capture conserved axes of immune readiness and dysregulation, which are predictive of future events and transfer across downstream tasks with lightweight predictors. We evaluate this hypothesis in two complementary settings of disease progression prediction and flu vaccine response prediction.

Rheumatoid arthritis (RA) is often preceded by an “at-risk” phase marked by anti-citrullinated protein antibodies (ACPA) in the absence of clinically apparent synovitis (i.e., clinical RA) [71, 72]. However, a substantial fraction (30%-70%) of ACPA^+^ individuals do not develop clinical RA over multi-year follow-up [73, 74]. Therefore, being able to precisely predict the imminent conversion in at-risk individuals will prompt more preemptive intervention, which could prevent or delay future tissue damage. We studied an ACPA^+^ at-risk cohort [75] longitudinally profiled with PBMC scRNA-seq (Figure 4a). In total, 46 participants were enrolled, 16 of whom progressed to clinical RA (converters) and 30 of whom remained disease-free (non-converters), yielding 96 profiled samples with a mean follow-up of 533 days. Serial pre-diagnostic timepoints were captured for converters, while a single baseline profile was included for non-converters. To rigorously assess PULSAR’s ability to identify pre-symptomatic rheumatoid arthritis, we implement donor-level cross-validation that preserves the independence of individual patients (Figure 4b). A zero-shot PULSAR embedding coupled to an XGBoost [76] classifier was compared with Pseudobulk, scPoli [49], TGSig [70] (which was developed as gene signatures for predicting vaccine responsiveness and autoimmune disease activity), and a non-transcriptomic clinical-variable baseline comprising multiple patient demographical data and relevant clinical measurements, including `DAS28-CRP`as a clinically deployed disease activity score for established RA (Methods 4.3.3). During XGBoost training, the model learns from all available pre-diagnostic visits from donors in the training set. For evaluation on held-out donors, we tested two clinically relevant scenarios: (i) predicting progression from any pre-diagnostic timepoint by evaluating each visit independently, and (ii) the more challenging task of predicting future RA development using only the earliest available pre-diagnostic visit. We quantified performance with repeated stratified cross-validation (5 repeats across 3 folds). For all pre-diagnostic visits, PULSAR achieved the highest Accuracy and F1 among all comparators (Accuracy=0.7196, F1=0.7616), outperforming the second-best baseline pseudobulk (Accuracy=0.6790, F1=0.7127) and outperforming TGSig, scPoli, and the clinical-variable model. Specifically, PULSAR outperforms the clinical-variable baseline by 17.8% in Accuracy and 21.7% in F1 score. When evaluating the earliest pre-diagnostic visit, PULSAR retained the top rank (Accuracy=0.6787, F1=0.5921), maintaining a margin over pseudobulk (Accuracy=0.6692, F1=0.5381) and especially over the clinical-variable baseline, with 17.4% improvement in Accuracy and 51.4% improvement in F1 score (Figure 4c, Supplementary Figure S13).

**Figure 4.**
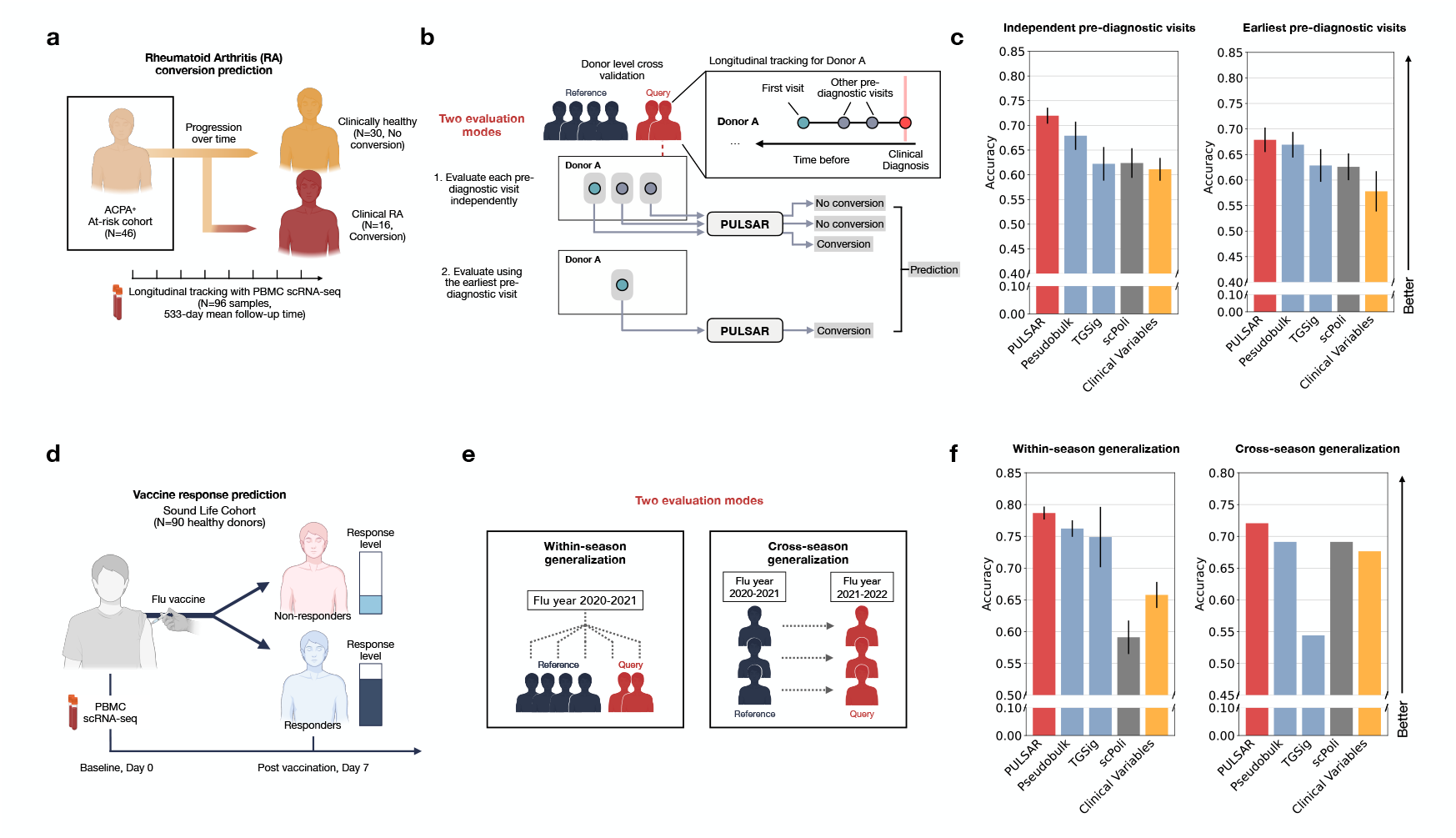
PULSAR improves prediction from baseline immune profile. **(a)** Schematic of the anti-citrullinated protein antibodies (ACPA^+^) at-risk cohort used for Rheumatoid Arthritis (RA) conversion prediction (46 participants; 96 PBMC profiles; mean follow-up 533 days), with outcomes of non-conversion (*n* = 30) or progression to clinical RA (*n* = 16). **(b)** Donor-level cross-validation strategy with two evaluation modes. Top: longitudinal tracking where each pre-diagnostic visit from the same donor is evaluated independently. Bottom: earliest visit only, testing the scenario of first-opportunity prediction. **(c)** RA conversion prediction performance comparing PULSAR to baseline methods. Accuracy scores for independent visits (left) and earliest visits (right). PULSAR is compared against pseudo-bulk, scPoli [49], TGSig [70], and clinical variables. Bar plots report mean *±* s.e.m. across repeated stratified cross-validation (5 repeats *×* 3 folds; *n* = 15 runs). **(e)** Schematic of the influenza vaccination cohort (Sound Life; 90 healthy donors) with baseline (day 0) PBMC scRNA-seq and day-7 hemagglutination inhibition assay outcome. **(f)** Two evaluation modes for vaccine response: within-season (train and test on different set of donors within the same flu year) and cross-season (train on one year, test on the subsequent year). **(g)** Vaccine response prediction benchmark reporting accuracy scores under two evaluation settings on within-season (left) and cross-season scenarios (right). Bar plots show mean *±* s.e.m. across repeated stratified cross-validation (5 repeats *×* 3 folds; *n* = 15 runs).

In our second case study, we examine inter-individual variability in responses to influenza vaccination, which reflects substantial heterogeneity in the human immune system arising from complex interactions between genetics and environment [5, 70]. Predicting these vaccine responses from baseline immune status is of broad interest, informing not only vaccine design but also our understanding of responses to other immune perturbations, such as infection and immunotherapy. We use the Sound Life cohort [42], which profiles *n* = 90 participants across multiple timepoints for two consecutive annual influenza vaccination seasons (Figure 4e). Specifically, we use the PBMC scRNA-seq sample on the day of receiving the vaccine to predict the vaccine responsiveness 7 days post-vaccination (Methods 4.3.4), defined by functional assay readout (Hemagglutination Inhibi-tion (HAI) assay, Supplementary Figure S12). We first ask whether baseline PULSAR embedding can predict vaccine responsiveness for held-out donors within a flu season. In our repeated stratified cross-validation in the 2020-2021 flu year, PULSAR achieved the best performance across metrics (Accuracy= 0.7867, F1= 0.8784). This exceeded the strongest baseline, pseudobulk (Accuracy=0.7622, F1=0.8626), and outperformed TGSig, scPoli. PULSAR further improved Accuracy by 19.6% and F1 by 16.1% from a clinical-variable model composed of clinical lab information and donor demographics. We also evaluated year-to-year generalisation, training on one flu season (2020-2021) and testing on the subsequent season (2021-2022). Despite antigenic drift and cohort effects, PULSAR again ranked first, with Accuracy=0.7206 and F1=0.8348. Pseudobulk trailed (Accuracy=0.6912, F1=0.8174) and TGSig lagged (Accuracy=0.5441 and F1=0.6353) (Figure 4f, Supplementary Figure S14). Performance that persists across seasons suggests that PULSAR encodes season-invariant axes of immune readiness, supporting robust and sample-efficient prediction of vaccine response from a single baseline draw.

### 2.5 PULSAR Simulates Cytokine Perturbation Across Physical Scales

A model that seamlessly navigates from molecules to cells to patients, simulating how perturbation impacts across biological scales, could help in therapeutic development and precision medicine. PULSAR achieves perturbation prediction at cross-scale resolutions through its hierarchical architecture, which learns representations that can be encoded and decoded across physical layers.

We apply PULSAR to model cytokine perturbation, addressing one of immunology’s most pressing clinical challenges [77]. As orchestrators of immune function that wire cell-cell communications, cytokines are key therapeutic targets across inflammatory, infectious, and malignant diseases [78–81], yet their effects are profoundly context-dependent, varying with cell type, dosage, and donor background.

To systematically evaluate cytokine response prediction, we analyze a comprehensive perturbation atlas comprising 10 million PBMCs from 12 donors [82]. Each donor’s cells are stimulated ex-vivo with 90 different cytokines plus controls, generating 1,080 paired baseline and post-perturbation single-cell profiles (Figure 5a). We test generalization across three challenging extrapolation scenarios: (i) novel cytokines not seen during training (Unseen cytokine), (ii) new donors with distinct immune backgrounds (Unseen donor), and (iii) unexplored donor-cytokine combinations (Unseen (donor+cytokine)).

**Figure 5.**
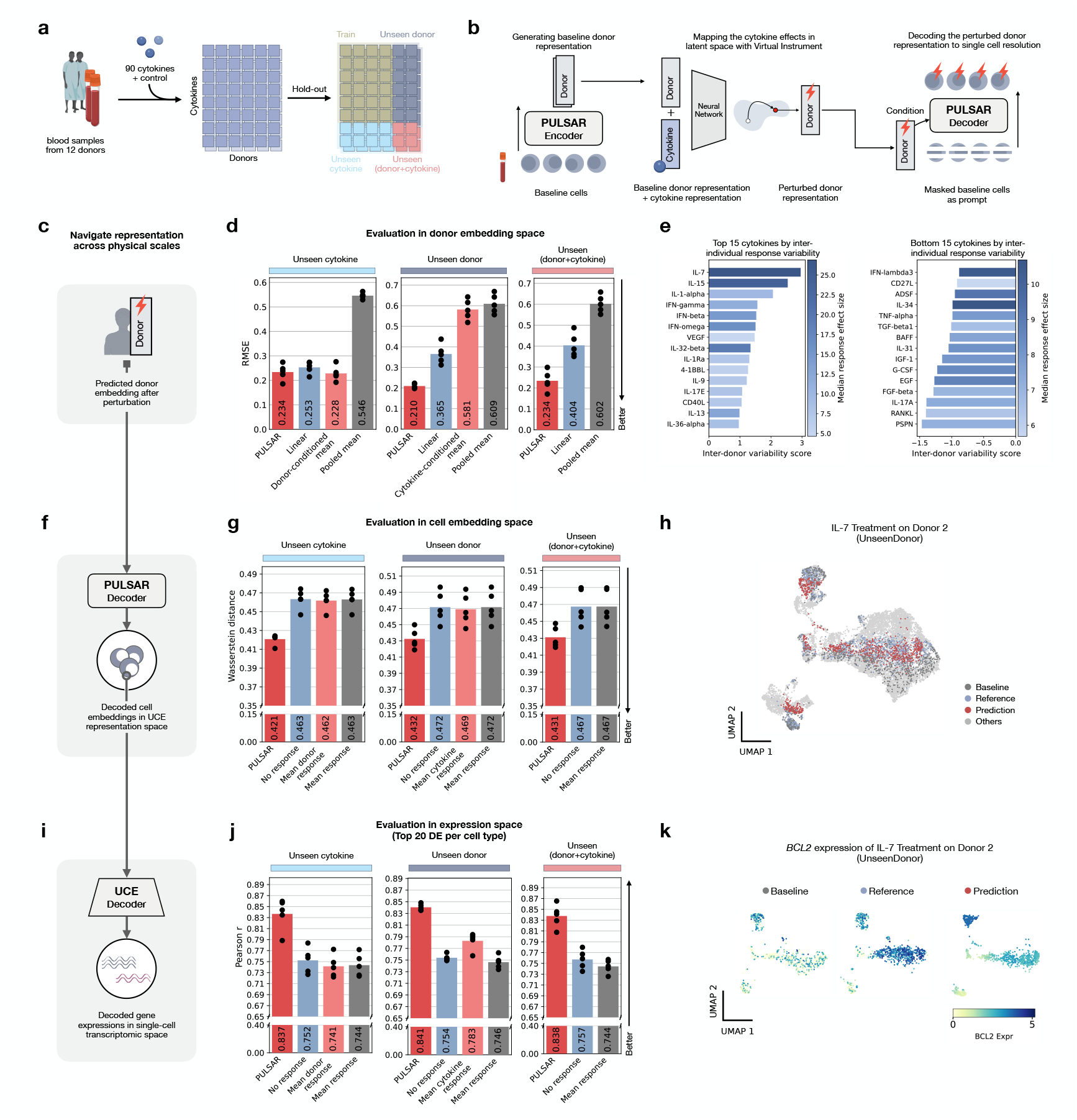
PULSAR simulates cytokine effects across physical scales. **(a)** Schematic overview of the cytokine perturbation benchmark evaluation: blood samples from twelve donors were stimulated with 90 distinct cytokines (plus a PBS control), and model performance is assessed by unseen donors, unseen cytokines, or unseen donor–cytokine combinations. **(b)** PULSAR perturbation modeling pipeline: a baseline donor embedding is transformed by a cytokine-conditioned neural “Virtual Instrument” to a perturbed donor embedding and decoded to single-cell embeddings. **(c, d)** Donor-space (or multicellular-space) evaluation. Reconstruction error (Root Mean Squared Error, RMSE; lower is better) comparing predicted to measured post-perturbation donor embeddings on held-out test cases. PULSAR is benchmarked against a linear regression operator and three mean baselines: a *donor-conditioned mean* (average perturbed embedding for the same donor across training cytokines), a *cytokine-conditioned mean* (average perturbed embedding for the same cytokine across training donors), and a *pooled mean* that ignores donor and cytokine identity. Each point represents a different test split (*n* = 5 runs in total). **(e)** Cytokines ranked by top-15 or bottom-15 inter-donor variability of response in donor space, with median effect size shown by colour. **(f, g)** Cellular embedding–space evaluation. Wasserstein distance is reported between predicted and measured post-perturbation cell-embedding distributions (lower is better) on held-out test cases. PULSAR is compared against a *no-response* baseline (cells unchanged from baseline) and mean-response heuristics: a *mean donor response* (average shift for that donor across training cytokines), a *mean cytokine response* (average shift for that cytokine across training donors), and a *pooled mean response* (global average shift). Each point represents a different test split (*n* = 5 runs in total). **(h)** UMAP of cell embeddings of IL-7 samples. Donor 2 cells are colored by prediction, reference and baseline. **(i, j)** Molecular space (decoded gene expression in single-cell RNA space) evaluation. Pearson correlation is reported between predicted and measured log-fold changes for the top 20 differentially expressed genes per cell type (higher is better) on held-out test cases. PULSAR is compared against the same no-response and mean-response baselines as in (f, g), now applied in expression space. Each point represents a different test split (*n* = 5 runs in total). **(k)** IL-7 case study on predicted *BCL2* gene expression for Donor 2. *BCL2* expression value is log-normalized.

PULSAR predicts these perturbation effects by traversing biological scales (Figure 5b,c,f,i). We first train a neural “Virtual Instrument” [1] that learns to map a donor’s baseline embedding to a post-perturbation state, conditioned on the cytokine’s protein embedding. To resolve cellular reprogramming, we employ conditional mask reconstructions: baseline cells are heavily masked and then reconstructed, prompted by the perturbed donor context. Finally, expression decoders project these predicted post-perturbation cell embeddings into gene expression space to resolve specific transcriptional programs (Methods 4.3.5). This hierarchical framework enables simultaneous reasoning across donor, cellular, and molecular resolutions. We benchmarked this capability by systematically withholding 4 donors and 20 cytokines across five random splits.

At the donor (or multicellular representation) scale, we assessed how closely each method reconstructs the post-perturbation donor state by computing the Root Mean Squared Error (RMSE) between predicted and measured donor embeddings. We benchmarked PULSAR against a linear instrument baseline and three mean-based heuristics: (1) assuming a donor has the same response to different cytokines (Donor-conditioned mean), (2) assuming a cytokine has the same effect on different donors (Cytokine-conditioned mean), and (3) assuming all treatments yield the same response (Pooled mean). PULSAR outperformed all baselines in the unseen-donor and unseen-donor-plus-cytokine regimes (Figure 5c,d). Most notably, for unseen donors, PULSAR reduced RMSE significantly, improving on the linear model by 42.5% and on the cytokine-conditioned mean by 63.9%. This confirms the model captures individualized context beyond generic averages. In the unseen-cytokine regime, however, PULSAR performed comparably to the donor-conditioned mean (0.234 vs. 0.228), indicating that when the donor is known, their baseline state provides a strong prior that dominates the prediction of novel stimuli.

Since cytokine responses vary significantly between individuals, we investigate whether PULSAR’s embeddings capture this heterogeneity. We compute an inter-donor variability score for each cytokine, quantifying how divergent donor trajectories are in response to the same stimulus (Figure 5e, all 90 targets at Supplementary Figure S15). IL-7 and IL-15 ranked highest, reflecting known individual differences in lymphocyte homeostasis [83, 84], while interferons (IFN-*γ*, IFN-*β*, IFN-*ω*) also showed high dispersion consistent with variable ISG induction [85]. In contrast, growth factors (e.g., G-CSF, EGF, FGF-*β*) elicited uniform responses across donors, suggesting these pathways are more conserved in peripheral blood under assay conditions.

At the cellular scale, we quantified how well PULSAR recapitulates held-out experiments by computing the Wasserstein distance between predicted and observed cell embedding distributions. PULSAR-generated cell embeddings consistently achieved the lowest distances across all testing regimes (unseen cytokine, unseen donor, and unseen both), outperforming four strong mean-based baselines (Figure 5g). These metrics indicate that PULSAR captures specific lineage-resolved shifts that generic population averages miss. Qualitatively, this is confirmed by the visualization of an IL-7 response in an unseen donor (Figure 5h): predicted cells (red) co-localize with measured reference cells (blue) in UMAP space, demonstrating a coherent functional shift away from the baseline state (grey).

At the molecular scale, we computed Pearson correlations for the top 20 differentially expressed genes across major cell types. PULSAR consistently outperformed mean-based heuristics across all generalization regimes, achieving correlations of 0.837-0.841 compared to 0.741-0.783 for baselines (Figure 5j). These results demonstrate that PULSAR reconstructs cell-type-specific transcriptional programs rather than merely shifting expression profiles. This accuracy is exemplified by the IL-7 response in an unseen donor (Figure 5k), where the model correctly predicts *BCL2* upregulation in targeted clusters, mirroring the canonical survival signaling pathway [86].

### 2.6 PULSAR Interpretability Reveals Cell-Type-Specific Disease and Severity Signatures

While predicting donor disease status and responses to perturbation is useful, the black-box nature of deep learning models poses a major challenge for explaining and understanding the underlying biological mechanisms. PULSAR’s multiscale hierarchical architecture, which links genes, cells, and donors in a single model, is naturally suited for interpretability because it preserves structure at each physical layer and aggregates it into a donor-level representation. We therefore ask whether the learned signals in the Multicellular Transformer of PULSAR could be used as a discovery tool to reveal cell-type-specific disease drivers and clinically relevant immune dynamics.

To find the cellular drivers of PULSAR’s donor representations, we perform a pan-disease attention-based analysis using our curated cross-disease atlas (`DONOR×EMBED`). We first standardize cell type labels across 2,804 PBMC samples (~10 million single cells) using a simple cell type classifier trained on UCE embeddings, and extract each cell’s attention weights from the Multicellular Transformer (Figure 6a, Methods 4.3.6). Aggregating these per-cell scores by major immune cell subset reveals disease-specific attention patterns (Figure 6b) that recapitulate known immunopathology. For example, autoimmune disorders, such as systemic lupus erythematosus (SLE), show a preferential weighting of memory B cells and antibody-secreting plasma cells. CD4^+^ effector-memory T cells in RA, reflecting autoantibody production and chronic T-cell help. Viral infections (COVID-19, influenza) are marked by elevated attention to innate effectors, including NK cells and NK-recruiting subsets, mirroring early antiviral responses, whereas chronic viral infection (HIV) shifts weight toward regulatory and central-memory T cells, in line with immune exhaustion and homeostatic proliferation.

**Figure 6.**
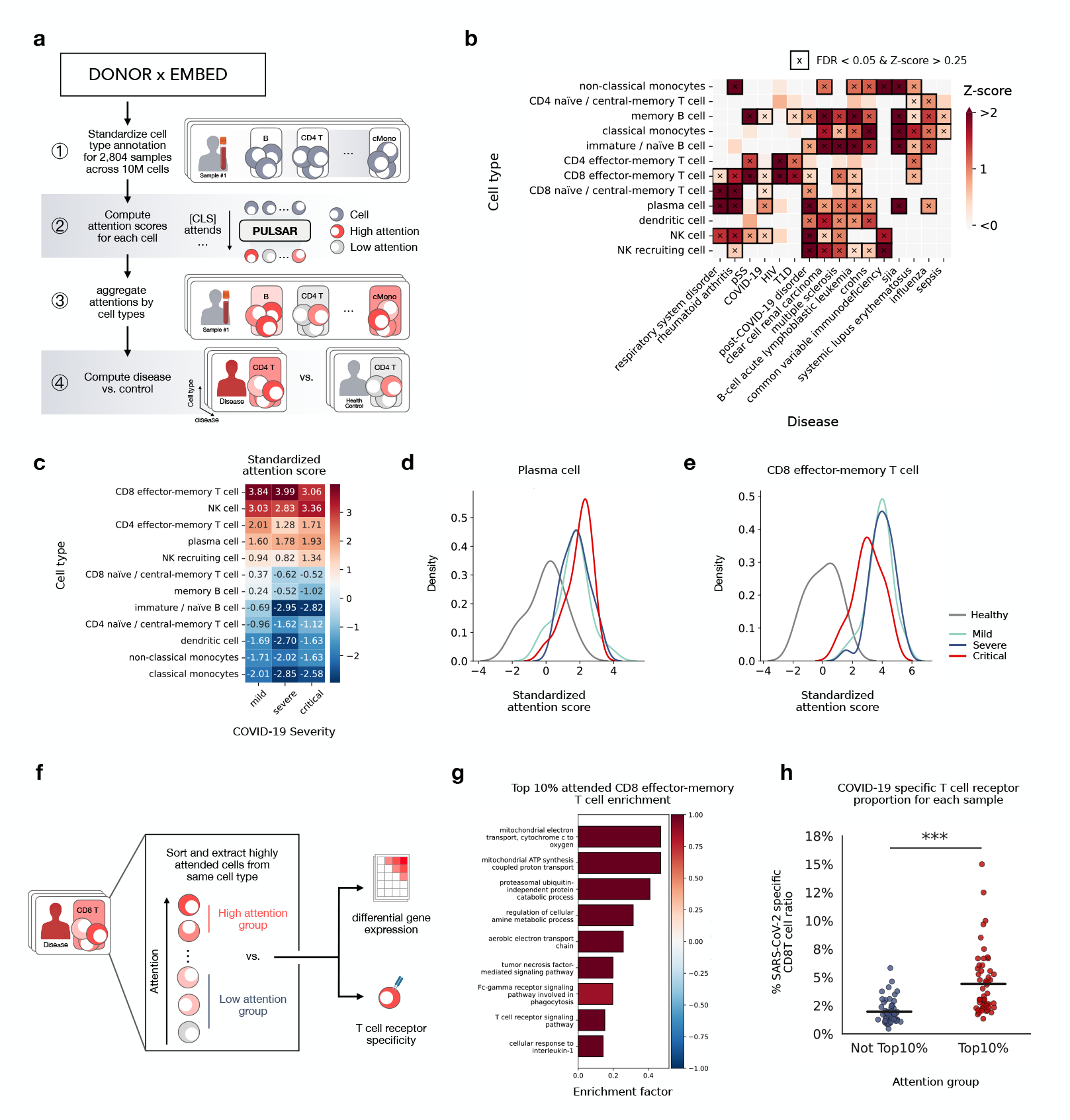
Attention analysis of PULSAR reveals cell-type–specific disease and severity signatures. **(a)** Schematic of the PULSAR attention analysis pipeline: single-cell annotations are standardized across 2,804 samples (~ 10 M cells) in `DONOR×EMBED`collection, per-cell attention scores are computed by the multicellular Transformer’s `[CLS]`token, and scores are aggregated by cell type to compare disease versus control. **(b)** Heatmap of standardized attention z-scores for major immune subsets across disease cohorts in `DONOR×EMBED`. Grids marked with “*×*” denote FDR *<* 0.05 and Z-score *>* 0.25. **(c)** COVID-19 severity stratification of attention: heatmap of Z-scores in key cell types for mild, severe and critical patients. **(d**,**e)** Kernel density estimates of standardized attention scores in plasma cells (d) and CD8 effector-memory T cells (e) across healthy (grey), mild (teal), severe (blue) and critical (red) COVID-19 samples. **(f)** Isolation of top and bottom-attention cells within the same cell type for downstream differential gene expression analysis and TCR specificity analysis. **(g)** Bar plot of the top 10 enriched GO pathways in the top 10% attended CD8 effector-memory T cells in COVID-19 samples. **(h)** Proportion of SARS-CoV-2-specific CD8 T cells in high versus low attention groups (*n* = 54, ^***^*p <* 0.001, two-sided test). Each point represents a sample, and the bar indicates the mean percentage (Not Top10% mean: 1.97%, Top10% mean: 4.41%).

Next, we analyze the annotated COMBAT cohort [87] to investigate whether PULSAR’s at-tention dynamics track clinical progression in COVID-19. Stratifying patients by severity (mild, severe, and critical), we observe progressive shifts in cell-type-specific attention that mirrored established pathology (Figure 6c). Plasma cell attention increases monotonically from mild (*Z* = 1.60) to critical disease (*Z* = 1.93), consistent with the escalating humoral response and hypergammaglobulinemia characteristic of severe infection [88]. Likewise, cytotoxic populations, specifically CD8 effector-memory T cells and NK cells, show peak attention in severe and critical patients (*Z* = 3.99 and *Z* = 3.36, respectively), reflecting their activated roles in viral clearance and immunopathology. Kernel density distributions confirm these trends (Figure 6d,e), revealing distinct positive shifts in attention distributions corresponding to worsening disease severity.

To further link attention to molecular function, we sort cells of the same type into top-versus bottom-attention groups (Top 10% highly attended cells vs. the rest, Figure 6f) and conducted differential gene expression (Supplementary Figure S16) for COVID-19 patient samples. In CD8 effector-memory T cells, the top 10% highest-attention cells are enriched for genes involved in mitochondrial electron transport, proteasome-mediated catabolism, and T cell receptor signaling against the rest cells (Figure 6g). Based on this signature, we hypothesize that the model picks up metabolically active, antigen-engaged T cells in COVID samples.

We then test whether high attention among T cells correlates with antigen specificity using paired TCR-sequencing data [89]. By mapping single-cell TCR sequences to a TCR antigen specificity database [90] (Methods 4.3.6), we find that high-attention T cells harbor a significantly larger fraction of virus-specific clonotypes compared to lowly attended counterparts (mean 4.41% vs. 1.97%, *p <* 0.001, two-sided test; Figure 6h). PULSAR thus identifies antigen-specific T cells solely from transcriptomic data, without TCR sequence information. This demonstrates that attention weights provide biological interpretability, identifying disease-relevant cells and revealing functional programs that could inform therapeutic targeting.

## 3 Discussion

We present PULSAR, a multi-scale and multicellular foundation model that links molecular, cellular, and donor-level biology into a unified predictive and generative framework. Applied to human peripheral immunity, PULSAR converts single-cell transcriptomes into donor embeddings that support disease classification, biomarker and clinical event prediction, and generative cytokine perturbation modeling while maintaining interpretability.

PULSAR’s capabilities open new avenues for the application of scRNA-seq in precision medicine [13]. Current clinical trials often rely on coarse stratification by age, sex, and disease severity scores that fail to capture the immunological heterogeneity driving treatment responses. PULSAR enables molecular patient stratification. For instance, tracking longitudinal risk for Rheumatoid Arthritis currently requires multi-year efforts with high attrition. Accurately quantifying risk using PULSAR could enrich trials for likely converters, reducing cohort sizes and accelerating readouts [91]. This principle extends to therapy selection as well. Because PULSAR prospectively predicts individualized cytokine responses, it can identify likely responders to immunomodulators and optimize dosing to achieve desired pathway engagement. This could ultimately support responder-enrichment designs and early futility decisions grounded in single-cell biology.

Despite these advances, several limitations and open questions remain. First, PULSAR is currently trained on dissociated single-cell data, which strips cells of their spatial context. While this works well for liquid tissues like PBMCs, spatial transcriptomics will be crucial for investigating the multicellular biology of solid tissues. Foundation model architectures must be developed specifically to incorporate tissue architecture and cell-cell communication dependent on physical proximity. Second, while scRNA-seq provides a rich view of cellular transcriptomic states, it captures only a single modality and currently ignores immune receptor repertoires, clonal expansion, and individual genetic features that shape transcriptional response over a lifetime. Integrat-ing paired TCR and BCR sequencing, along with genetic variants that modulate transcriptional responses, would allow PULSAR’s molecular layer to encode donor-specific immune histories, including imprinting from latent viral infections and other environmental exposures (e.g. EBV or CMV) [92, 93]. Third, while our hierarchical architecture enables bottom-up reasoning, fully back-propagating gradients across physical scales to refine representations remains both a computational and data challenge.

Overall, PULSAR shows that a multi-scale and multicellular foundation model can bridge molecular, cellular, and multicellular biology to support clinically relevant inference at the individual level. As omics measurements are increasingly deployed in the clinical setting, we expect PULSAR to become a core component of patient-specific digital twins and data-driven precision medicine.

## 4 Methods

### 4.1 PULSAR Architecture

PULSAR architecture employs a hierarchical encoder-decoder design following the multi-scale principles of the AI Virtual Cell (AIVC) [1]. Throughout this work, we use “sample representation,” “multicellular representation,” and “donor representation” interchangeably.

Formally, let 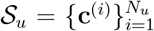 denote the set of single cell profiled from sample *u*, where *N*_*u*_ is the number of cells in sample *u*. Cell **c**^(*i*)^ is described by its corresponding expression profile **x**^(*i*)^ ℕ^*G*^ across *G* genes. PULSAR Encoder is defined by `PULSAREncoder`: 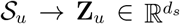 that maps unordered cell sets to fixed *d*_*s*_-dimensional sample embeddings (**Z**_*u*_) for sample *u*. PULSAR is also paired with a PULSAR Decoder `PULSARDecoder`: 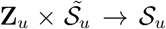 that reconstructs cellular information conditioned on sample embeddings and masked cells, where 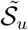 stands for the masked cells.

PULSAR processes information through three distinct physical scales: Molecular, Cellular, and Multicellular. The model has three core components: (1) cell set sampling, (2) multi-scale representations, and (3) multicellular transformer.

#### 4.1.1 Cell Set Sampling

Per-sample cell counts could exhibit high variability due to variations in sequencing assays and sample preparation protocols. To enable efficient training while preserving the cellular heterogeneity in each sample, we implement a cell sampling strategy. For each training tissue sample, we uniformly sample *L* cells per sample (default *L* = 1, 024, taking the nearest exponent of 2 from the median per-sample cell count of the Onek1k dataset [43] at 1246). Formally, for a sample *u*, we denote its full cell set by

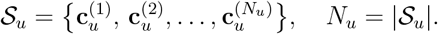

We wish to obtain exactly *L* cells (default *L* = 1, 024) as

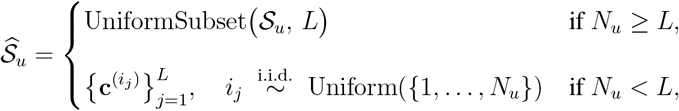

where UniformSubset(𝒮_*u*_, *L*) draws a size-*L* subset without replacement, uniformly over all 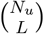 choices, and when *N*_*u*_ *< L* we sample with replacement so that 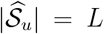 is always maintained.

#### 4.1.2 PULSAR Uses Multi-scale Representation

PULSAR processes biological information hierarchically across three scales: molecular, cellular, and multicellular. We tokenize each biological unit and integrate these representations across scales, leveraging Transformers. At the molecular level, we use ESM2 (`esm2_t48_15B_UR50D`) [19] to encode protein sequences from expressed genes, capturing evolutionary and biophysical properties of individual proteins. At the cellular level, we employ Universal Cell Embeddings (UCE, 33 Layers 650M Parameters) [21] to transform sparse single-cell expression profiles into dense, batch-corrected representations that capture cellular states by incorporating molecular-level ESM protein embeddings. These high-quality pre-trained embeddings from established foundation models enable PULSAR’s multicellular transformer to focus on learning emergent multicellular patterns at the tissue level.

##### Molecular Scale: Gene Representation

Each expressed gene *g* in cell **c**^(*i*)^ is denoted by 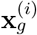. *g* represents any protein-coding gene. We extract the corresponding gene embedding (denoted as 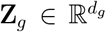) using average ESM2 protein embedding for each protein product from gene *g* [19]. Specifically for ESM2, it yields a *d*_*g*_-dimensional gene embedding with *d*_*g*_ = 5, 120.

##### Cellular Scale: Cell Representation

Single-cell expression profiles are first transformed into a cell embedding space before aggregating into a multicellular representation. We use Universal Cell Embedding (UCE) to aggregate the molecular embeddings into a unified cell embedding with minimal batch effects [21]. UCE leverages gene molecular representations from ESM2 to generate a cellular embedding based on the abundance of gene expression. UCE samples 1, 024 gene tokens, with replacement, from each cell’s expressed gene set, where the sampling probability for gene *g* is weighted by its log-normalized expression. UCE then groups the gene embeddings according to their chromosome positions and generates a cell-level embedding 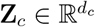 through a Transformer backbone. In the context of UCE, *d*_*c*_ is fixed to 1,280.

##### Multicellular Scale: Sample Representation

For each sample 𝒮_*u*_, we construct the cell set 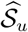 through sampling to a fixed number of cells (Section 4.1.1). We then transform this cell set into the UCE cell embedding space and feed the bag of cells into the Multicellular Transformer model (PULSAR Encoder) for a multicellular sample-level representation (**Z**_*u*_) at fixed dimension size *d*_*s*_. By default, PULSAR generates an embedding size at *d*_*s*_ = 512.

#### 4.1.3 Multicellular Transformer

Multicellular Transformers are used both in PULSAR Encoder and PULSAR Decoder to model the multicellular data. The core challenge in learning multicellular representations for dissociated samples lies in capturing two fundamental aspects of tissue organization: (1) cellular composition with the diversity and abundance of cell types present and coordinated intercellular patterns (the functional relationships among cells) and (2) single-cell state alternation, which involves the dynamic states and transitions of individual cells within the multicellular context. Traditional approaches that treat cells independently fail to model these emergent multicellular properties that arise from collective cellular behavior [23–25].

To address this challenge, we tokenize cellular states and employ a transformer backbone [45, 95] with a multi-head self-attention mechanism to learn coordinated intercellular patterns in a permutation-invariant manner. With cells as tokens, the transformer model captures specific transcriptomic alterations at single-cell resolution while maintaining a global field of view covering the entire cell set. Furthermore, the Multicellular Transformer is trained in a cell-embedding space, which mitigates batch effects and enables learning across diverse sample conditions.

##### Input Embedding

Formally, we represent the set of selected cell embedding 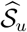 as 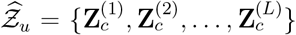, where each 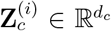 represents the UCE cell embedding of cell *i*. Cell embeddings are projected to the hidden dimension of the model (*d*_*h*_) with a MLP 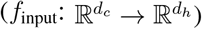. We represent each projected cell embedding as 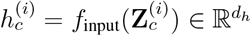 for each cell *i*. We construct the input sequence with an additional `[CLS]`token:

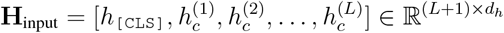

where 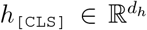 is a learnable token designed to aggregate the information of the sequence.

##### Transformer Encoding

The input embeddings are then fed through *N* Transformer layers for contextualizing the sample embedding.

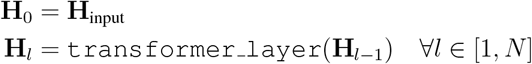

After *N* layers, we extract the final `[CLS]`token 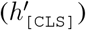 from **H**_*N*_ and project it to the sample embedding space with learnable projection 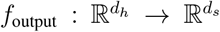. For sample 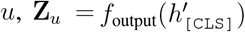 is final sample-level representation.

Each `TransformerLayer`consists of:

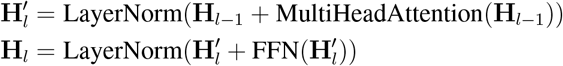

where MultiHeadAttention enables each cell embedding to attend to all other cells in the sample in a bi-directional way, and FFN is a two-layer feed-forward network with GELU activation [96].

#### Transformer Decoding

Another transformer model is used in PULSAR Decoder during the pretraining task to reconstruct the masked cell embeddings and during the perturbation setting to generate post-perturbation cell embeddings, conditioned on a sample-level representation. The training details are discussed in (Methods 4.2.2), and the architecture is described below:

First, some masked cell embeddings are projected to the hidden dimension through the masked sample input MLP 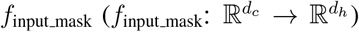, constructing 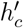. A sample representation **Z**_*u*_ is used as a condition token to construct the input sequence as:

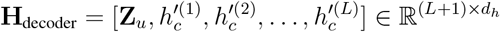

The decoder employs *M* transformer layers with attention mechanisms to distribute sample-level information into cell-level predictions:

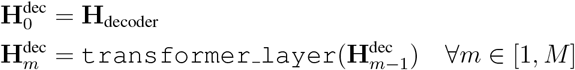

The transformers here use bidirectional attention without causal mask. The final cell embeddings are extracted from 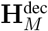 (excluding the donor token) and projected back to the cell embedding space through 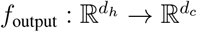:

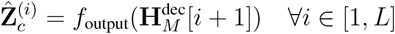

### 4.2 PULSAR Pretraining

#### 4.2.1 Construction of Pretraining Corpus

We assembled a large-scale single-cell RNA sequencing corpus for PULSAR pretraining by leveraging the CZ CELLxGENE Census database (version `LTS 2023-07-25`) [29, 46]. The initial data collection yielded 36,227,903 unique single-cell transcriptomes after removing duplicate entries. Each cell profile was accompanied by standardized metadata annotations and precomputed UCE representations. To resolve sample labels from individual cells’ metadata, we used a grouping strategy based on three key metadata fields: `dataset id`, `donor id`, and `tissue`. Each unique combination of these identifiers is labeled as a distinct sample based on the assumption that cells sharing these three attributes represent a coherent biological unit from a single individual, experimental batch, and anatomical location. This grouping strategy identified 7,034 unique sample combinations across the corpus. Quality control is performed by filtering samples with fewer than 200 cells, resulting in a total of 6,807 samples for Stage 1 training. At this stage, we have gathered data from 36,207,954 cells across 53 tissues and 69 disease conditions. This ensures the model can learn unbiased, high-quality sample-level representation across the full spectrum of human tissues and conditions.

For the Stage 2 pretraining (Continual pretraining), we created a specialized subset that focuses exclusively on peripheral blood samples to enhance the model’s capacity for immunological profiling. From the Stage 1 corpus, we extracted all samples annotated as blood tissue (by filtering `tissue_general`), applying identical quality control criteria. This yielded 8,735,936 cells distributed across 2,588 high-quality blood samples, providing dense coverage of immune cell diversity and donor-specific variation.

#### 4.2.2 Pretraining Task: Masked Cell Modeling

To pretrain PULSAR in a self-supervised manner, we developed Masked Cell Modeling, a generative pretraining task that learns to encode multicellular systems at single-cell resolution. The PULSAR Encoder generates a multicellular embedding from a visible subset of cells, while the PULSAR Decoder reconstructs the remaining cells from this embedding.

During pretraining, each sampled cell set 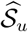 of length *L* is split into two subsets: a visible set 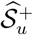 of size *L*^+^ and a reconstruction target 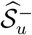 of size *L*^−^. The visible set 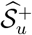 is processed by the PULSAR Encoder to compute the sample embedding, while 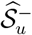 serves as the reconstruction target for the PULSAR Decoder.

For reconstruction, we mask dimensions in the UCE embedding space, replace masked elements with learnable elements, and train the PULSAR Decoder to reconstruct these masked dimensions conditioned on the sample embedding.

To enhance learning effectiveness, we make three key design choices inspired by He et al. [97]: (1) Asymmetric architecture: The PULSAR Encoder uses *N* = 8 transformer layers while the PULSAR Decoder uses *M* = 6 layers, promoting stronger encoding capacity. (2) High invisible cell set ratio: We use a 15:85 visible-to-invisible split (i.e., *L*^+^ : *L*^−^ = 15% : 85%), requiring the sample embedding to extrapolate to unseen cell states. (3) Correlated masking: This adaptive masking strategy computes a masking prior from the covariance matrix of the UCE embedding space, performs hierarchical clustering to identify groups of correlated dimensions, then randomly masks 85% of these groups. We compute the covariance matrix empirically from OneK1K [43]. We apply correlated masking only during Stage 2 continual pretraining; for Stage 1 pretraining, we use random masking.

Formally, let 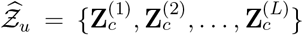 denote the UCE cell embeddings for sample *u* computed from 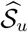, where each 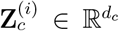. We randomly partition the cell indices [1, *L*] into visible and masked subsets such that |`visible`| = *L*^+^ = 0.15*L* and |`masked`| = *L*^−^ = 0.85*L*.

The visible cells 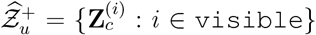 are encoded to generate the sample embedding:

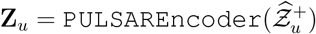

##### Random masking

For Stage 1 pretraining, we employ random dimension masking. For each masked cell 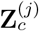 where *j* ∈ `masked`, we randomly select *p* proportion of the dimensions to mask. This generates a binary mask 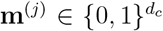 where each dimension *d* has probability *p* of being masked 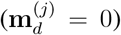 and probability (1 − *p*) of remaining visible 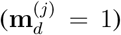. The masked cell embedding is constructed as:

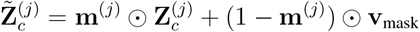

where 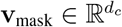 is a learnable mask token where **v**_mask_ = *v*_mask_ · **1** is a learnable constant vector (i.e., all elements equal to the same scalar *v*_mask_), and ⊙ denotes element-wise multiplication. During PULSAR training, we have *p* = 0.85.

##### Correlated masking

For Stage 2 continual pretraining, we use a more sophisticated masking strategy that exploits correlation structure in the UCE embedding space. Let 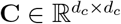 denote the covariance matrix computed across all cell embeddings in the training set. We perform hierarchical clustering on **C** to obtain *K* groups of correlated dimensions: 𝒢 = {*G*_1_, *G*_2_, …, *G*_*K*_} where 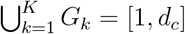 and *G*_*i*_ ∩ *G*_*j*_ = ∅ for *i* ≠ *j*.

During each training iteration, we randomly select *p × K* groups to mask, creating a binary mask 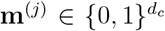 where 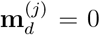 if dimension *d* belongs to a masked group, and 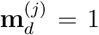 otherwise. The masked cell embedding is constructed identically to random masking using the same shared mask token **v**_mask_. PULSAR uses *p* = 0.85.

##### Decoding and training objective

The decoder reconstructs the original cell embeddings conditioned on the sample embedding:

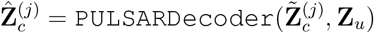

The pretraining objective minimizes the reconstruction loss only on masked dimensions:

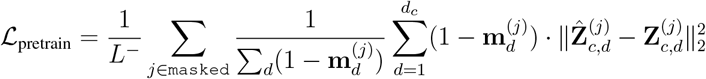

where the loss is computed only over masked dimensions for each cell.

##### Training details

The model was trained at `bf16`-mixed precision with AdamW (*β*_1_ = 0.9, *β*_2_ = 0.999, *ϵ* = 1 × 10^−8^) [98] with 2 A100 80GB GPUs using Distributed Data Parallel (DDP) for each stage. Training proceeded for 2,048 epochs with a learning rate of 5 *×* 10^−5^ and a global batch size of 192, with a 50 warm-up steps, cosine learning rate scheduler, and gradient clipping (max gradient norm at 1.0). During the training process, the model learns from over 19.7 billion total tokens.

### 4.3 Task Description

#### 4.3.1 Reference Mapping Task

To build a reference-based inflammation classification system, we assembled a comprehensive database of 2,804 scRNA-seq profiles with corresponding disease labels from 41 sources: 32 datasets from CellxGene [29, 46], 1 from the Single Cell Portal (SCP) [99], and 8 from additional curated sources. For external validation, we created a held-out test cohort of 254 profiles from 6 independent datasets (1 from CellxGene, 1 from SCP, and 4 from other sources). These test samples share the same disease categories as the reference but differ in key technical and biological variables, including sequencing platforms, patient demographics, and age distributions, ensuring robust evaluation of generalization performance. A complete list of datasets and publications is provided in Extended Data Table 1.

We standardized all expression data by aligning each cell’s gene expression vector to CellxGene’s 60,664-gene feature panel. Disease labels were harmonized using the Mondo Disease Ontology [100] following CellxGene schema, while inflammation categories were assigned using the classification framework following Jiménez-Gracia et al. [48].

For efficient reference-based classification, we pre-computed donor embeddings by sampling 1,024 cells per donor and processing them through PULSAR to generate 512-dimensional donor representations. These embeddings were then indexed in a vector database to enable rapid *k* = 3 nearest-neighbor queries for incoming samples using the Minkowski distance, followed by a majority vote to generate the final prediction. Baseline methods are evaluated in the same reference mapping setting.

To align PULSAR’s embedding space with disease labels, we further performed contrastive learning on the full model weights by training an additional 3-layer MLP projection head (PULSAR-aligned). The contrastive learning [101] framework constructs positive pairs from samples sharing the same disease label and negative pairs from samples with different diseases. We optimized the model using a margin-based contrastive loss function defined as:

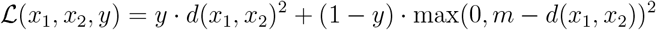

where *x*_1_ and *x*_2_ are the embedding vectors of the paired samples, *y* ∈ {0, 1} is the label indicating whether the pair shares the same disease (*y* = 1 for positive pairs, *y* = 0 for negative pairs), *d*(·, ·) denotes the Euclidean distance between embeddings, and *m* is the margin hyperparameter set to 0.65. This loss function minimizes the distance between positive pairs while enforcing a minimum separation of *m* between negative pairs. Training proceeded for 10 epochs using the Adam optimizer [102] with a learning rate of 5 *×* 10^−4^ and a batch size of 32, with the best model selected based on the validation loss.

Pseudobulk baselines were computed by averaging gene expression counts across all cells per donor, applying *L*1 normalization to standardize the library size, and followed by an `log1p` transformation. For the scPoli baseline, we followed the two-phase training protocol from the original publication [49]. In the “Reference building” phase, scPoli was trained on all samples from the CellxGene database. In the subsequent “Reference mapping” phase, we loaded the pretrained weights and fine-tuned the model to learn sample-specific representations (i.e., “learned conditional embeddings”) for both the `DONOR×EMBED`cohort and the external validation cohort. We adopted the hyperparameters from their `PBMC atlas`model configuration: a 3-layer encoder with 128 hidden units, a 30-dimensional latent space, a 20-dimensional embedding space, *η* = 1, and alpha epoch annealing of 1,000 epochs, except we expanded the input features to align the gene list in CellxGene. The cell type composition baseline is a 30-dimensional vector that describes the abundance of the 30 different cell types. The cell types across different datasets are re-mapped using UCE embedding with a pretrained logistic classifier. A detailed list of annotated cell types is included in the Supplementary Table S1.

#### 4.3.2 Protein Expression Regression Task

To evaluate the PULSAR embeddings’ performance on reflecting system-level biomarkers, we leverage a dataset with scRNA-seq profiles paired with Olink proteomic panels [54]. We use the PULSAR model to generate zero-shot donor embeddings for each sample using 1,024 cells. We then utilize a Ridge regression head for predicting each individual protein. During the training process, only the projection head is trained while the PULSAR transformer backbone remains frozen. Baselines are set up similarly to reference mapping (Method 4.3.1), computing pseudo-bulk profiles from the top 2,000 highly variable genes computed with Scanpy [103] and `seurat`flavour.

To visualize the protein co-expression network and contextualize PULSAR’s prediction performance, we constructed a protein-protein interaction network where edges represent co-expression patterns in ground-truth data and nodes encode prediction quality. We computed pairwise Spearman correlations between all 454 proteins using measured plasma protein levels across samples. Protein pairs with |*ρ*| ≥ 0.3 were retained as edges, with edge widths scaled by correlation strength. For each protein (node), we quantified PULSAR’s prediction accuracy by computing the Spearman correlation between predicted and measured levels across all samples, followed by FDR correction using the Benjamini-Hochberg procedure. Node colors represent prediction correlation (*ρ* ∈ [0,1]), and node sizes encode statistical significance as −log_10_(FDR). The network was visualized using a spring layout algorithm (NetworkX [104] implementation with optimal distance between nodes *k* = 0.25).

#### 4.3.3 RA Conversion Prediction Task

To evaluate PULSAR embedding performance in predicting rheumatoid arthritis (RA) conversion, we used a longitudinal scRNA-seq dataset profiling an at-risk cohort [75]. We generated zero-shot donor embeddings for each sample using randomly sampled 512 cells and trained an XGBoost classifier [76] to predict binary conversion outcomes. The design decisions are based on limited sample sizes. For comparison, we implemented three transcriptome-based baseline methods: scPoli was fine-tuned using the reference mapping approach (Methods 4.3.1). Pseudobulk profiles were computed from the top 1,200 highly variable genes. For the TGSig baseline [5, 70], we subset pseudo-bulk profiles to the ten-gene TGSig signature (*C15orf57* (also known as *CCDC32*), *LONP2, PAPSS2, EPHB1, ADAM12, SMC1A, RETN, ENPP1, CD101* and *C2orf63* (also known as *CLHC1*)) to train the classifier. We also constructed a clinical variable baseline using seven donor-specific clinical measurements at blood draw: `anti-cyclic citrullinated peptide-3 (CCP3)`, `C-Reactive Protein (CRP)`, `Sex`, `BMI`, `DAS28-CRP`, `rheumatoid factor IgA`and `rheumatoid factor IgM`. To construct evaluation protocols, we excluded diagnostic visit timepoints for all donors with multiple visits. One converter had only one sample at the diagnostic visit and was included for training to retain sample size.

#### 4.3.4 Vaccine Response Prediction Task

To evaluate PULSAR embedding performance in predicting influenza vaccine responsiveness, we used the Sound Life cohort [42], which longitudinally tracked 90 participants across two flu seasons (2020-2021 and 2021-2022). We used baseline PBMC scRNA-seq profiles (vaccination day, day 0) to predict binary response labels derived from Meso Scale Discovery (MSD) hemagglutination inhibition (HAI) assays on day 7 post-vaccination. The HAI assay quantifies antibody-mediated inhibition of hemagglutination in paired serum samples, with higher inhibition indicating stronger functional antibody activity. For 2020-2021 (flu year 1), HAI tested three strains: `A/Guangdong`, `B/Phuket`and `B/Washington`. For 2021-2022 (flu year 2), HAI tested `A/Cambodia`, `B/Phuket`and `B/Washington`. Response was measured as normalized percentage inhibition, with a positive response defined by above 30% inhibition for each strain according to the manufacturer’s recommendation. [42]. We then aggregated strain-specific responses to donor-level responses using a one-of-any criterion (responder if any strain exceeded the threshold). We generated zero-shot donor embeddings for each day-0 sample using 1,024 randomly sampled cells and trained a logistic regression classifier to predict binary response labels. As with RA conversion prediction (Methods 4.3.3), we implemented baseline methods similarly, except clinical variables included 8 variables: `absolute lymphocyte count`, `absolute neutrophil count`, `absolute monocyte count`, `percentage of lymphocytes`, `white blood cell count`, `platelet count`, `age at first visit` and `sex`. We applied an additional class-weight-balancing to the TGSig, scPoli, and clinical-variable baselines to mitigate overfitting.

To benchmark cross-individual generalization, we used the 2020-2021 flu year (*n* = 90 participants) and constructed a repeated stratified cross-validation. For cross-season generalization, we trained on 2020-2021 data (*n* = 90) and tested on held-out 2021-2022 data (*n* = 68 same donors in the 2020-2021 flu year cohort).

#### 4.3.5 Cytokine Perturbation Prediction Task

We used the Parse-PBMC-10M dataset [82] for training and evaluation. To assess generalization, we randomly split the donors and cytokines to construct held-out test cases. To enable PULSAR to predict perturbation responses across three physical scales, our pipeline comprises three modules: (1) a Virtual Instrument [1], a generative model operating in donor embedding space, that transforms baseline donor embeddings to post-perturbation donor embeddings conditioned on cytokine identity, (2) a PULSAR decoder that maps baseline cell embeddings to post-perturbation cell embeddings conditioned on the predicted post-perturbation donor embedding, and (3) a UCE embedding-to-expression decoder that enables exploration across physical scales.

The training procedure consists of two stages. First, we finetuned the PULSAR Encoder and PULSAR Decoder on training donors and cytokines using context expansion, enabling a wider field of view for cell distribution, and an adjusted masking strategy optimized for generation tasks. This training step adapts PULSAR to the distribution of perturbation responses. Second, we trained both the Virtual Instrument and the UCE expression decoder in parallel using the finetuned PULSAR model’s embedding.

During inference, the pipeline operates sequentially: (1) the fine-tuned PULSAR Encoder generates the baseline donor embedding from sampled cells, (2) the Virtual Instrument predicts the post-perturbation donor embedding conditioned on cytokine identity, (3) the fine-tuned PULSAR Decoder generates post-perturbation cell embeddings conditioned on the predicted donor embed-ding, using highly masked baseline cells as prompts, and (4) optionally, the UCE embedding-to-expression decoder translates cell embeddings to gene expression profiles for downstream analysis.

##### Problem formulation and data split

First, we define the set of all donors and cytokines as 𝒟 and respectively, where |𝒟| = 12 and |𝒞| = 90. There is another PBS treatment control for each donor, which serves as the baseline state. We randomly split the training and test dataset following: |𝒟_train_| = 8 and |𝒟_test_| = 4; |𝒞_train_| = 70 and |𝒞_test_| = 20. For each donor and cytokine pair (*d, c*) ∈ 𝒟 *×* 𝒞, we define four test scenarios based on the training splits:

1. Training: (*d, c*) ∈ 𝒟_train_ *×* 𝒞_train_. Both donor and cytokine were seen during training (*n* = 560 pairs)
2. Donor extrapolation: (*d, c*) ∈ 𝒟_test_ *×* 𝒞_train_. Unseen donor, seen cytokine (*n* = 280 pairs)
3. Cytokine extrapolation: (*d, c*) ∈ 𝒟_train_ *×* 𝒞_test_. Seen donor, unseen cytokine (*n* = 160 pairs)
4. Full extrapolation: (*d, c*) ∈ 𝒟_test_ *×* 𝒞_test_. Unseen donor-and-cytokine during training (*n* = 80 pairs)

This partitioning enables systematic evaluation of the model’s generalization capabilities across different biological axes, with a total of |𝒟| *×* |𝒞| = 1080 donor-cytokine pairs. During evaluation, we repeat this random split 5 times.

Our objective is to learn a mapping function *f* : 𝒳_baseline_ *×* 𝒞 → 𝒳_perturbed_ that predicts the postperturbation state given a baseline state and cytokine identity. Critically, the state representation 𝒳 can be expressed across three hierarchical physical scales: (1) donor-level space 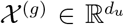, capturing population-level responses, (2) cell embedding space 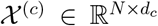 where *N* is the number of cells and *d*_*c*_ is the embedding dimension, and (3) gene expression space 𝒳^(*g*)^ ∈ ℝ^*N ×g*^ where *g* is the number of genes. Our modular pipeline decomposes this mapping into scale-specific transformations: 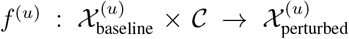 for the Virtual Instrument, 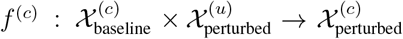 for the multicellular decoder, and 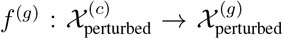 for expression decoding.

##### Virtual Instrument

PULSAR’s Virtual Instrument [1] simulates the cytokine perturbation effect in the donor embedding space, which learns a 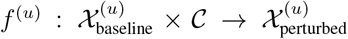. The vir-tual instrument employs a Feature-wise Linear Modulation (FiLM) architecture [105] to transform baseline donor embeddings (**x**_baseline_ ∈ ℝ^512^) into post-perturbation donor embeddings (**y** ∈ ℝ^512^) conditioned on perturbation context (**c** ∈ ℝ^5120^). For conditioning each cytokine (context), we use the cytokine’s ESM2 protein product embeddings. In the special case where the cytokine is a heterodimer, we apply the average of the two protein embeddings. This representation supports the generalization to unseen cytokines. The transformation proceeds with the donor embedding being processed through a donor-specific network:

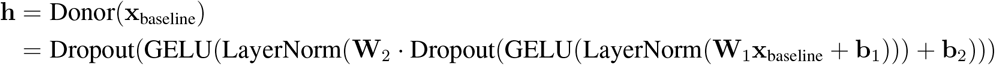

where **W**_1_, **W**_2_ ∈ ℝ^512*×*512^ are weight matrices, and **h** ∈ ℝ^512^ represents the processed donor features.

Then the perturbation context generates modulation parameters through parallel networks:

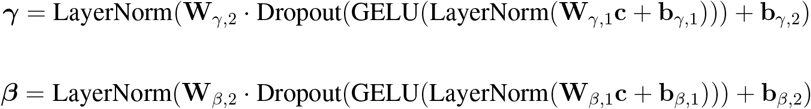

where **W**_*γ*,1_, **W**_*β*,1_ ∈ ℝ^1024*×*5120^, **W**_*γ*,2_, **W**_*β*,2_ ∈ ℝ^512*×*1024^, and ***γ, β*** ∈ ℝ^512^ are the scale and shift parameters, respectively.

FiLM modulation is applied element-wise:

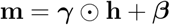

where ⊙ denotes element-wise multiplication. Finally, the modulated features are projected to the output space with a residual connection [106]:

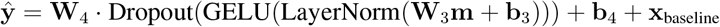

where **W**_3_, **W**_4_ ∈ ℝ^512*×*512^ and **ŷ** ∈ ℝ^512^ is the predicted post-perturbation donor embedding.

The model is trained by minimizing a composite loss function:

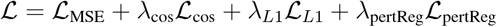

where:

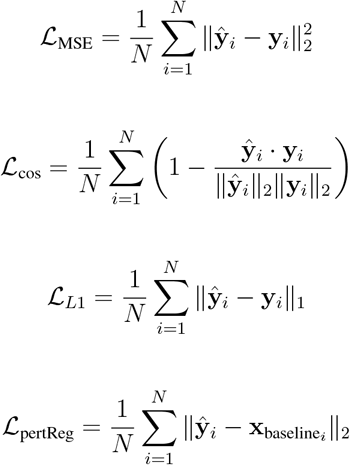

with *λ*_cos_ = 0.1, *λ*_*L*1_ = 0.05, and *λ*_pertReg_ = 0.01. We employed the Adam optimizer [102] with learning rate *η* = 0.001, weight decay 10^−5^, and gradient clipping (maximum norm = 0.5). The model was trained for up to 25,000 epochs, using a 90:10 train-validation split and early stopping (patience = 50 epochs).

##### Multicellular decoder for generating perturbed cell embeddings

We finetuned the model using Masked Cell Modelling (Methods 4.2.2) similar to pretraining on the training donors and cytokines. To maximize the generative capability, we performed context expansion by encoding 1,024 cells and decoding 1,024 cells at once, and further increased the random masking rate to 0.95. During inference, the decoder operates conditionally on the Virtual Instrument’s prediction. Given baseline cell embeddings 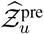 and the predicted post-perturbation donor embedding **ŷ**_*u*_, we first mask 95% of the baseline cells randomly. The decoder then reconstructs all 1,024 cell embeddings conditioned on both the unmasked cell (providing baseline cellular context) prompt and the donor embedding (encoding the perturbation effect). This approach enables the model to generate post-perturbation cell states that maintain consistency with the donor-level response while capturing cell-type-specific and stochastic variations in perturbation effects. We use the finetuned PULSAR Decoder to approximate 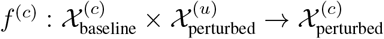. The predicted cell embeddings can be directly used for downstream analysis or further decoded to gene expression space using the UCE embedding-to-expression decoder.

##### UCE-expression decoder

To enable interpretation and analysis at the gene expression level 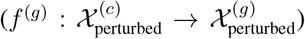, we trained a UCE-Expression Decoder that maps cell embeddings back to gene expression space. This decoder consists of a 2-layer MLP that transforms 1,280-dimensional UCE cell embeddings to expression values for 7,933 highly variable genes. The model uses a zero-inflated negative binomial (ZINB) reconstruction loss. We trained the decoder using the Adam optimizer with a learning rate of 0.005 and a batch size of 4,096 for 64 epochs.

##### Evaluation across physical scales

To assess model performance at each biological scale, we designed scale-specific evaluation metrics. At the donor (multicellular) scale, we computed the distance between predicted and measured donor embeddings. At the cellular scale, we used distributional metrics to compare generated versus reference cell-embedding distributions. At the molecular scale, we stratified predictions by cell type and evaluated cell-type-specific gene expression changes.

##### Donor-level evaluation

For each held-out donor–cytokine pair (*d, c*) ∈ (𝒟_test_, 𝒞_test_) we compare the predicted post-perturbation donor embedding **ŷ**_*d,c*_ ∈ ℝ^512^ with the measured embedding **y**_*d,c*_ using

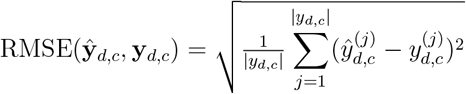

We compare PULSAR against four different baselines:

Linear regression: Concatenate the baseline donor embedding 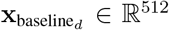 and the cy-tokine protein embedding **p**_*c*_ ∈ ℝ^*P*^, and fit a linear map:

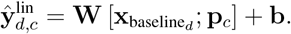

- Donor-conditioned mean (used in unseen-cytokine): average the training post-perturbation donor embeddings for the same donor:

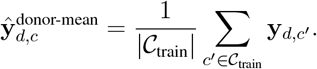
- Cytokine-conditioned mean (used in unseen-donor): average across training donors for the same cytokine:

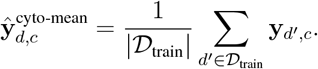
- Pooled mean: the global mean over all training pairs

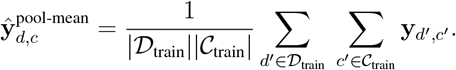

##### Cell-level evaluation

To quantify distributional agreement between predicted and measured post-perturbation single-cell states, we compute the Wasserstein–2 (W2) distance with exact optimal transport (OT). Let 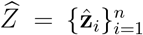 and 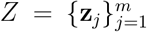 be the predicted and reference cell-embedding sets in UCE space. We use uniform masses 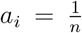 and 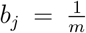, and a quadratic Euclidean ground cost 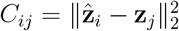. The OT objective is

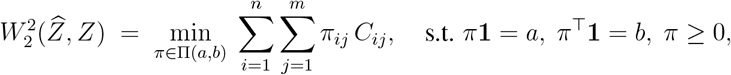

and we report 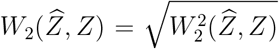. In implementation, we solve the exact OT problem via `ot.emd2`[107, 108].

We compare PULSAR against four baselines by constructing mean-response predictors by applying an average displacement vector to the baseline cells. Define the displacement for a training pair (*d, c*) as 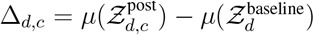, where 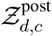 is the post reference perturabtion cell embedding for donor *d* after treatment 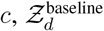 is the baseline cell embeddings for donor *d* and *µ*(·) is the sample mean in embedding space. Then:

- No response: 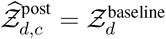.
- Mean donor response (used for unseen cytokine): use the donor specific average:

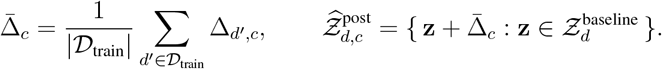
- Mean cytokine response (used for unseen donor):

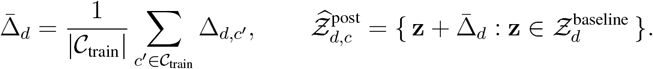
- Pooled mean: use the global average displacement:

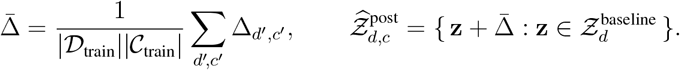

##### Expression-level evaluation

As inspired by existing work [109], for each cell type, we define a reference panel of the top *K* differentially expressed genes (here *K* = 20) from the measured post-versus pre-perturbation comparison. We then score predictions by the Pearson correlation between predicted and observed log-fold changes on this panel, reporting the mean across all major cell types (frequency ≥ 5%) and all held-out pairs. Cell types for generated cells are inferred using a UCE embedding-based logistic classifier, similar to Methods 4.3.1. Baselines mirror the cell-level benchmarks but are constructed in expression space rather than embedding space (i.e., using cytokine-, donor-, or globally averaged log-fold-change vectors).

##### Quantify cytokine individualization in donor space

For each donor *d* ∈ {1, …, *D*} = 𝒟 and cytokine *c* ∈ {1, …, *C*} = 𝒞, we define the displacement in donor-embedding space as 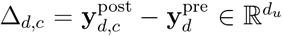. To reduce background noise, we whiten displacements by estimating a Ledoit-Wolf covariance matrix [110] and to decorrelate them and set unit variance for 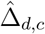. Directional concordance across donors is summarised by *R*_*c*_:

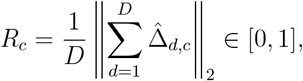

and we define a directional individualization score for cytokine *c* as 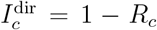 with higher values indicating more donor-divergent response directions. To reflect effect size, we aggregate the median magnitude medMag_*c*_ = median_*d*_(*m*_*d,c*_) with 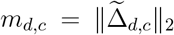. Finally, we form a composite ranking by z-scoring components and summing:

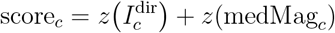

and sort cytokines by score_*c*_ in descending order. Cytokines with both large effect size and divergent donor trajectories rank highest.

#### 4.3.6 Interpretability Analysis

Model attention scores across donor samples are compared across cell types and within each cell type using the `DONOR×EMBED`cohort. Cell types in `DONOR×EMBED`were inferred using a pre-trained logistic regression classifier trained on reference cell type annotations, yielding 30 distinct cell type categories, similar to Methods 4.3.1. Each cell’s UCE embedding was mapped to its most probable cell type using this classifier and aggregated into a coarse cell type.

We extracted attention weights from disease-label-aligned PULSAR’s Multicellular Transformer layers, specifically focusing on the PULSAR Encoder `[CLS]`token’s attention to all other cell tokens, as `[CLS]`is supervised to capture the cell-level alterations. For a sample with *N* cells (including the `[CLS]`token at position 0), we extracted the attention matrix **A** ∈ ℝ^(*N* +1)*×*(*N* +1)^ from the specified layer (by default, last layer before final layer). The attention score for each cell *i* was computed as:

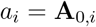

where *A*_0,*i*_ represents the attention scores from the `[CLS]`token to cell *i*.

To quantify attention patterns by cell type, we computed the average attention score for each cell type *c* within each donor *d*:

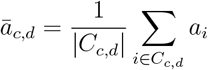

where *C*_*c,d*_ denotes the set of cells belonging to cell type *c* in donor *d*.

To account for background variations across datasets (e.g., technical artifacts and donor demographics) while preserving disease-specific signals, we employed a linear mixed model:

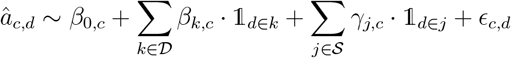

where 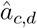 is the modeled attention for cell type *c* in donor *d*, represents the set of disease categories (with “normal” as the reference category), represents the set of dataset IDs, *β*_*k,c*_ captures the disease-specific effect for disease *k* relative to normal controls, *γ*_*j,c*_ captures the dataset-specific batch effect, and `𝟙`denotes indicator functions. The model is fitted using the observed values 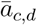 as the dependent variable.

To enable comparison across cell types with different baseline attention levels, we computed standardized effect sizes (z-scores) for each disease-cell type combination as 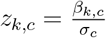 where *β*_*k,c*_ is the estimated disease effect for disease *k* in cell type *c* (relative to normal), and *σ*_*c*_ is the within-cell-type standard deviation of the observed attention scores 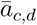 computed across all samples with degrees of freedom correction. Statistical significance was assessed using the regression model’s p-values for each disease coefficient, followed by Benjamini-Hochberg false discovery rate (FDR) correction across all cell type-disease combinations to account for multiple testing. Cell type-disease pairs with FDR *<* 0.05 and standardized effect size *z >* 0.25 were considered significantly enriched.

To enable the multi-omic analysis across attention groups, we utilize a dataset [89] with 125 donors paired with single-cell RNA-seq and paired single-cell TCR-seq. Antigen specificity of T-cell receptors (TCR) was inferred by referencing the single-cell TCR-seq against a curated reference (VDJdb [90]) using scirpy [111, 112]. CDR3 amino-acid sequences were then compared against VDJdb using a sequence-identity search to capture identical receptors. For each matched clonotype, the corresponding antigen species and epitope annotations were retained. We binarize each TCR’s COVID-19 specificity by the presence of `SARS-CoV-2`antigen species hit. We also filter out samples that do not show `SARS-CoV-2`antigen presence in either highly attended or lowly attended cell groups. This process yields 54 samples for final comparison.

## Data Availability

All analyzed datasets are publicly available. `DONOR×EMBED`can be accessed at HuggingfaceHub https://huggingface.co/datasets/KuanP/PULSAR_DONORxEMBED_aligned and https://huggingface.co/datasets/KuanP/PULSAR_DONORxEMBED_zero_shot for disease aligned and zero-shot PULSAR embeddings respectively. A full list of data used to construct `DONOR×EMBED`and the external validation cohort can be found in the Extended Data Table 1. Data used for predicting plasma proteomics can be downloaded at Array Express: E-MTAB-10129. Data used for Rheumatoid Arthritis conversion prediction can be downloaded at https://apps.allenimmunology.org/aifi/insights/ra-progression/. Sound Life cohort for vaccine response prediction can be accessed at https://apps.allenimmunology.org/aifi/insights/dynamics-imm-health-age/cohorts/. Data for COVID-19 severity analysis is accessed at https://cellxgene.cziscience.com/collections/8f126edf-5405-4731-8374-b5ce11f53e82. Data used in the COVID multi-omic analysis can be downloaded at Array Express: E-MTAB-9357.

## Code Availability

PULSAR is written in Python using Pytorch (Version 2.8.0) [113], Scanpy (Version 11.11.4) [103], Numpy (Version 2.3.3) [114], and HuggingFace Transformers (Version 4.57.0) [115] libraries. Linear and logistic regression models are trained using sklearn (Version 1.5.2) [116]. XG-Boost model is trained with xgboost package (Version 3.0.2) [76]. The source code and environment configuration are available on GitHub at https://github.com/snap-stanford/PULSAR/. Model weights are publicly available at HuggingfaceHub, zero-shot model: https://huggingface.co/KuanP/PULSAR-pbmc and disease-aligned model: https://huggingface.co/KuanP/PULSAR-aligned.

## Acknowledgements

We thank Zinaida Good, Anshul Kundaje, Tony Wyss-Coray, Xiao-jun Li, Eric Lubeck, Orit Rozenblatt Rosen, Rok Sosič, Kexin Huang, Hanchen Wang, Zoe Piran, Michael Bereket, Moritz Schaefer, Rita Li, Alan DenAdel, Haotian Cui, Kameron Rodrigues, Kristin Tsui, Can Ergen and all members of the SNAP group and OCTO group for discussions and for providing feedback on our manuscript. We also gratefully acknowledge the support of NSF under Nos. OAC-1835598 (CINES), CCF-1918940 (Expeditions), DMS-2327709 (IHBEM), IIS-2403318 (III); Stanford Data Applications Initiative, Wu Tsai Neurosciences Institute, Stanford Institute for Human-Centered AI, Chan Zuckerberg Initiative, Amazon, Genentech, and SAP. A.R. is supported by NIH/NCI R01CA296792. We thank the Allen Institute founder, P.G. Allen, for his vision, encouragement, and support. The content is solely the responsibility of the authors and does not necessarily represent the official views of the funding entities. Plots are created using BioRender.

## Author Information

K.P., Y.R. and J.L. conceived the study. K.P., Y.R. and J.L. performed research, contributed new analytical tools, designed algorithmic frameworks, analyzed data and wrote the manuscript. K.P., Y.R., and K.K performed experiments and developed the software. K.K., Z.H., A.R., C.G., G.H. provided and analyzed the pre-RA and Sound Life data. J.L. supervised the study. All authors edited the manuscript.

## Supplementary Information

**Figure S1.**
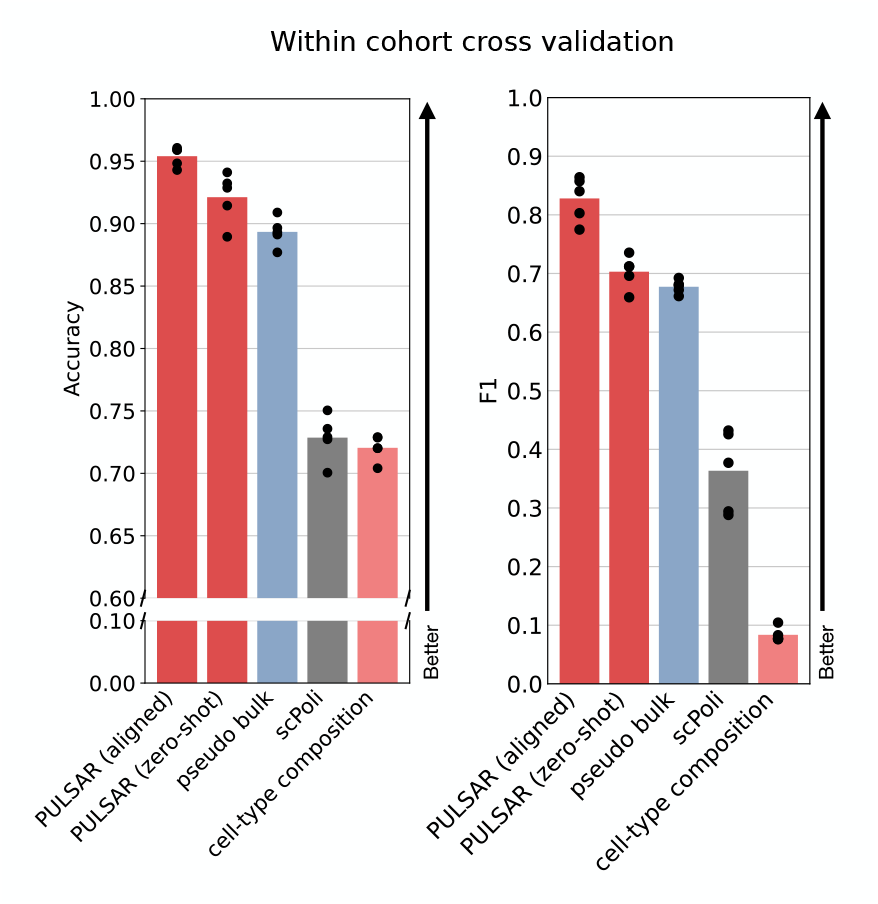
Performance benchmark for within-cohort cross-validation of disease classification. Accuracy (left) and F1 (right). Points denote the mean over 5-fold runs.

**Figure S2.**
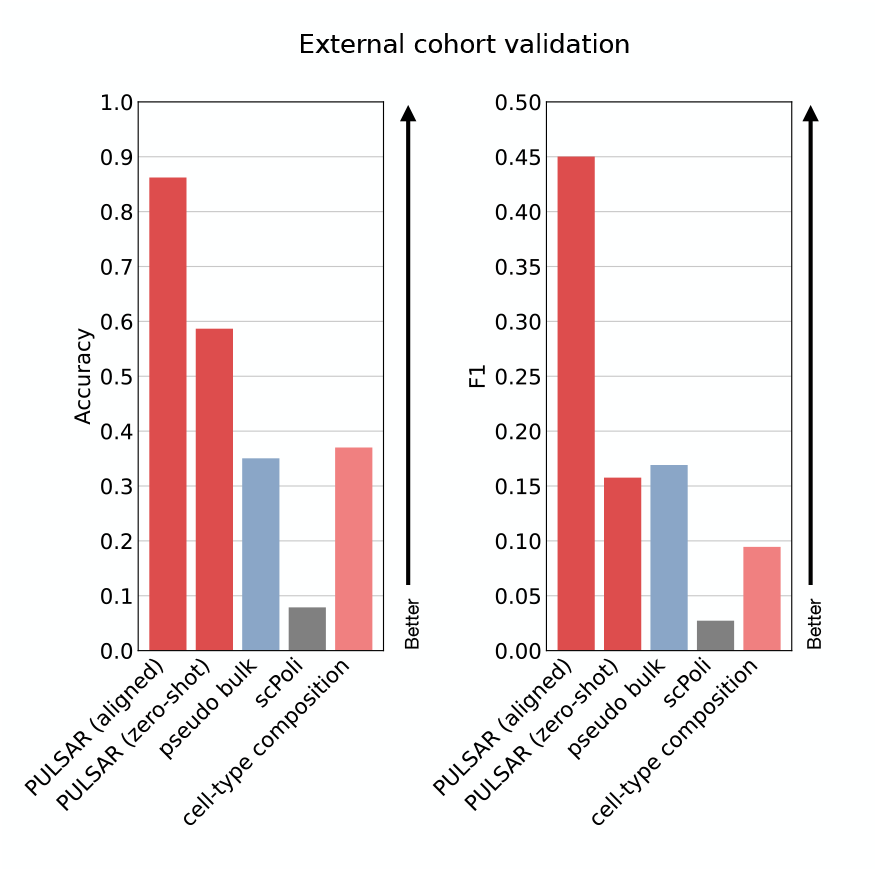
Performance benchmark for external cohort cross-validation of disease classification. Accuracy (left) and F1 (right). Points denote the mean over 5-fold runs.

**Figure S3.**
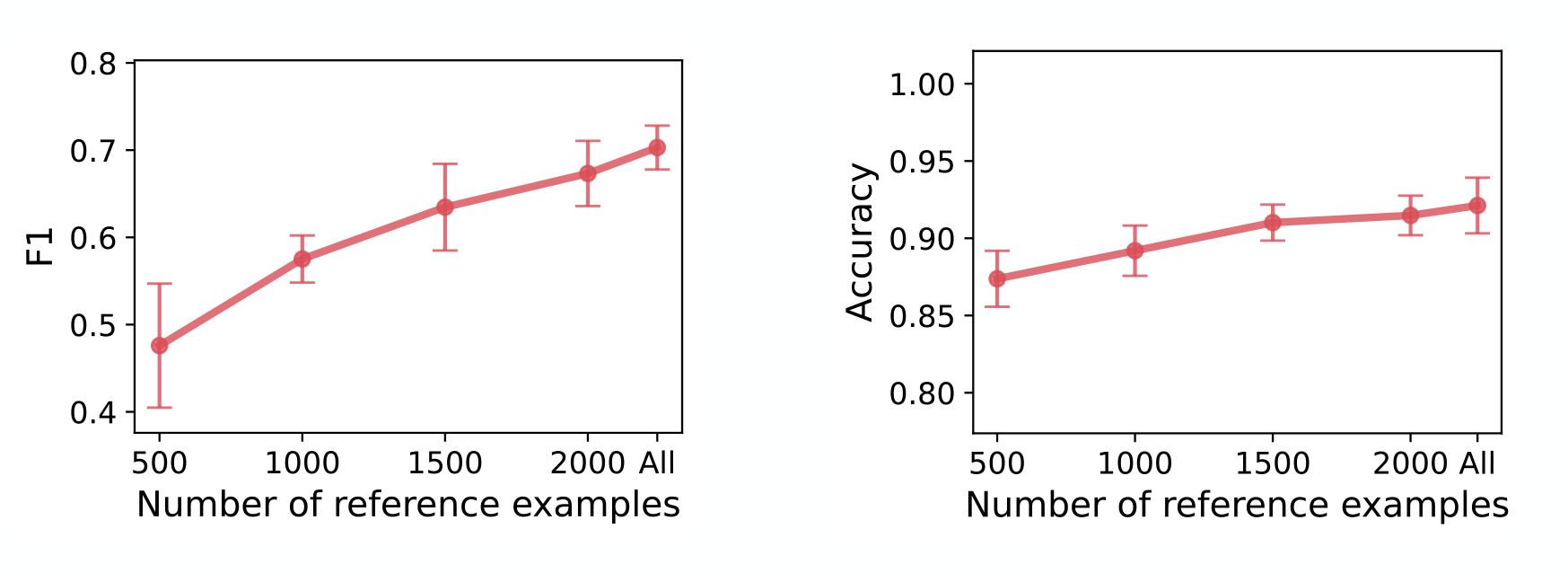
Performance scales with the size of the reference vector database. Macro F1 (left) and Accuracy (right) as a function of the number of reference examples retrieved from the vector database in the reference split. All is 2243 samples. Points denote the mean over 5-fold runs, error bars indicate s.e.m. We observed a scaling effect as the size of the reference vector database increased. Expanding from 500 to 1,500 reference examples yielded the largest gains (Macro F1 from 0.588 to 0.669; Accuracy 0.875 to 0.891), with steady improvements up to the full reference corpus (Macro F1 0.712; Accuracy 0.889). The trends indicate that expanding the reference corpus improves search accuracy.

**Figure S4.**
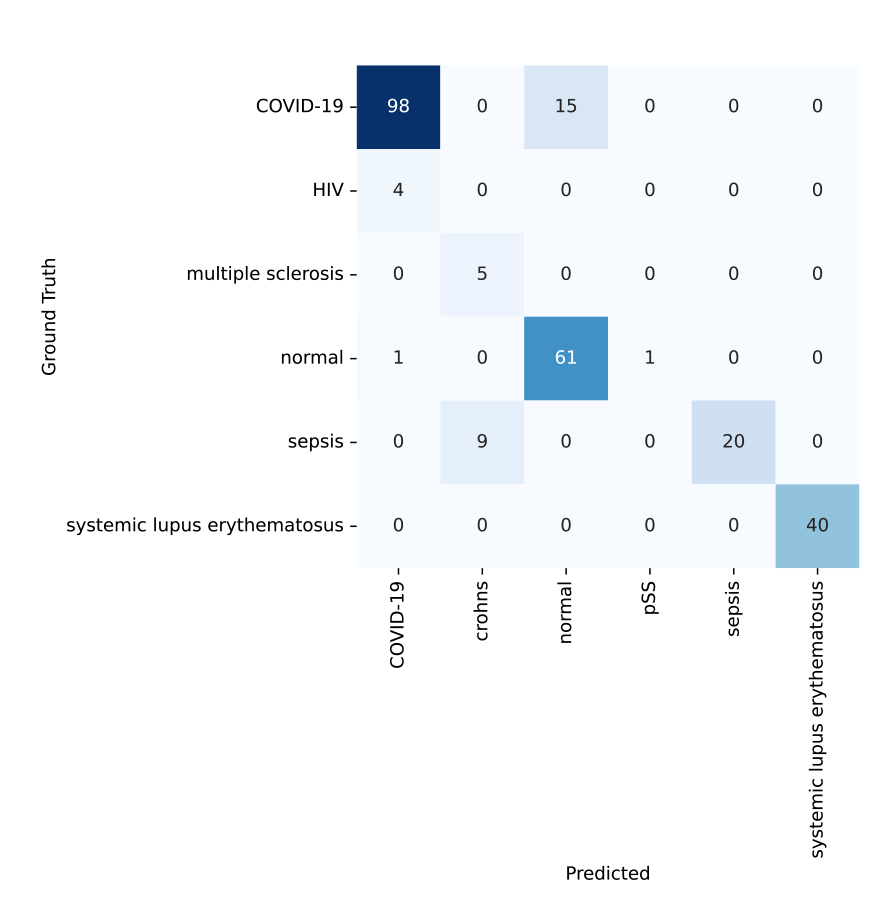
Confusion matrix on the external cohort. Confusion matrix summarizing predictions across an external cohort at the disease level.

**Figure S5.**
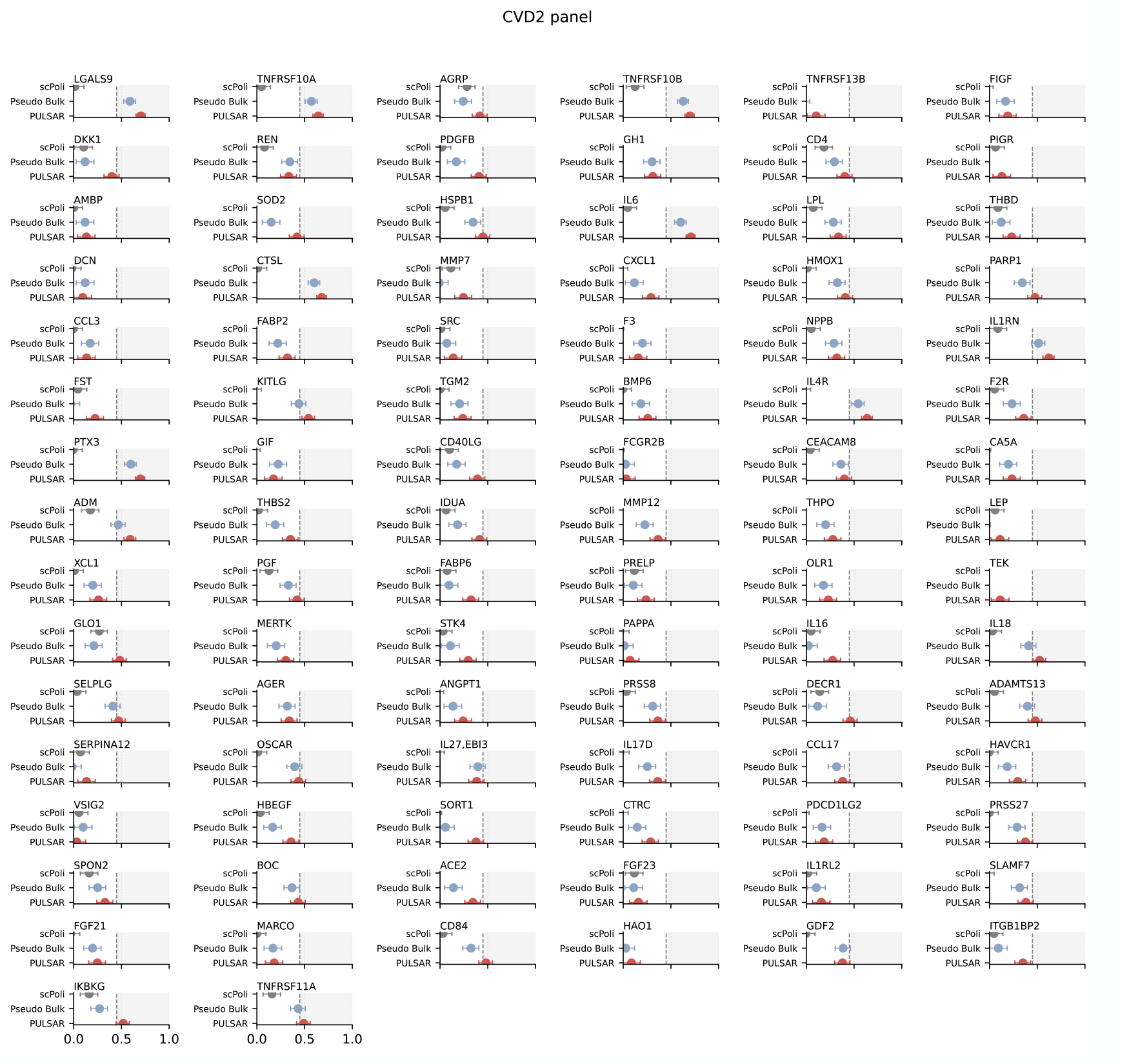
Protein prediction on the Olink CVD2 panel. Per-analyte performance comparing scPoli, Pseudo-bulk, and PULSAR for predicting plasma protein abundance from baseline PBMC profiles. Points denote the mean Spearman correlation between predicted and measured levels across cross-validation folds; horizontal bars indicate 95% CI. The vertical reference line marks the performance threshold at a Spearman correlation of 0.45.

**Figure S6.**
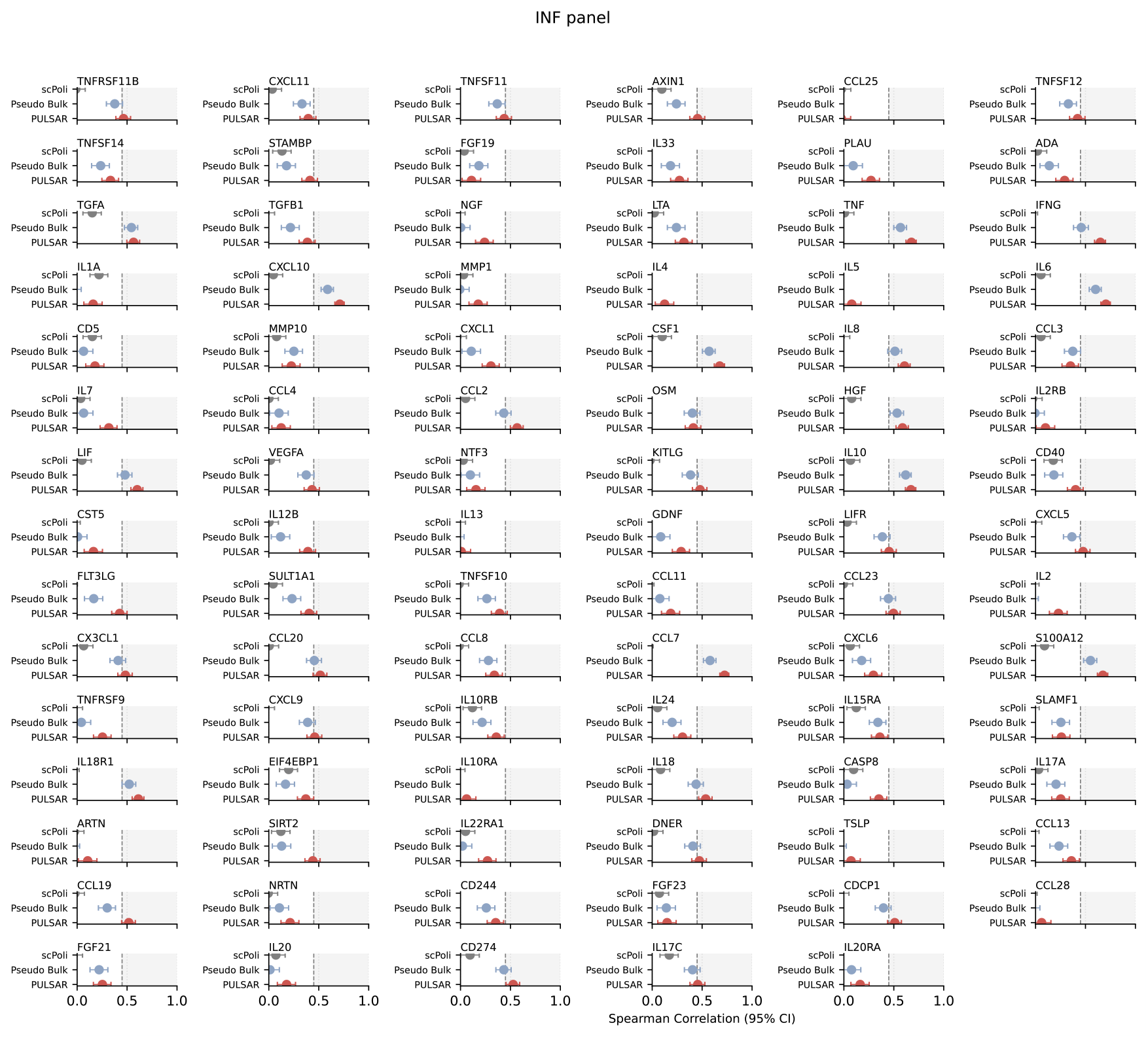
Protein prediction on the Olink INF panel. Per-analyte performance comparing scPoli, Pseudo-bulk, and PULSAR for predicting plasma protein abundance from baseline PBMC profiles. Points denote the mean Spearman correlation between predicted and measured levels across cross-validation folds; horizontal bars indicate 95% CI. The vertical reference line marks the performance threshold at a Spearman correlation of 0.45.

**Figure S7.**
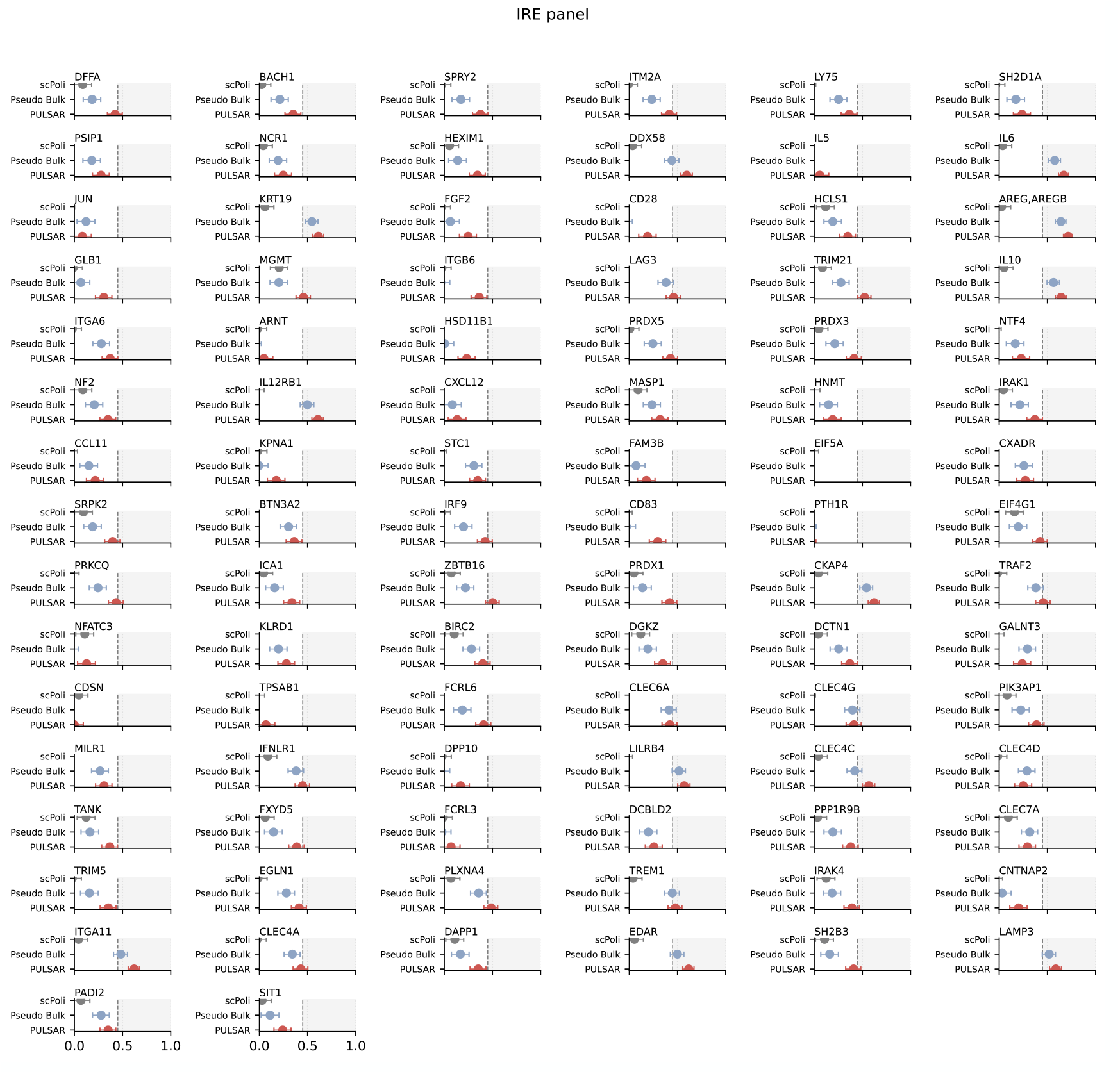
Protein prediction on the Olink IRE panel. Per-analyte performance comparing scPoli, Pseudo-bulk, and PULSAR for predicting plasma protein abundance from baseline PBMC profiles. Points denote the mean Spearman correlation between predicted and measured levels across cross-validation folds; horizontal bars indicate 95% CI. The vertical reference line marks the performance threshold at a Spearman correlation of 0.45.

**Figure S8.**
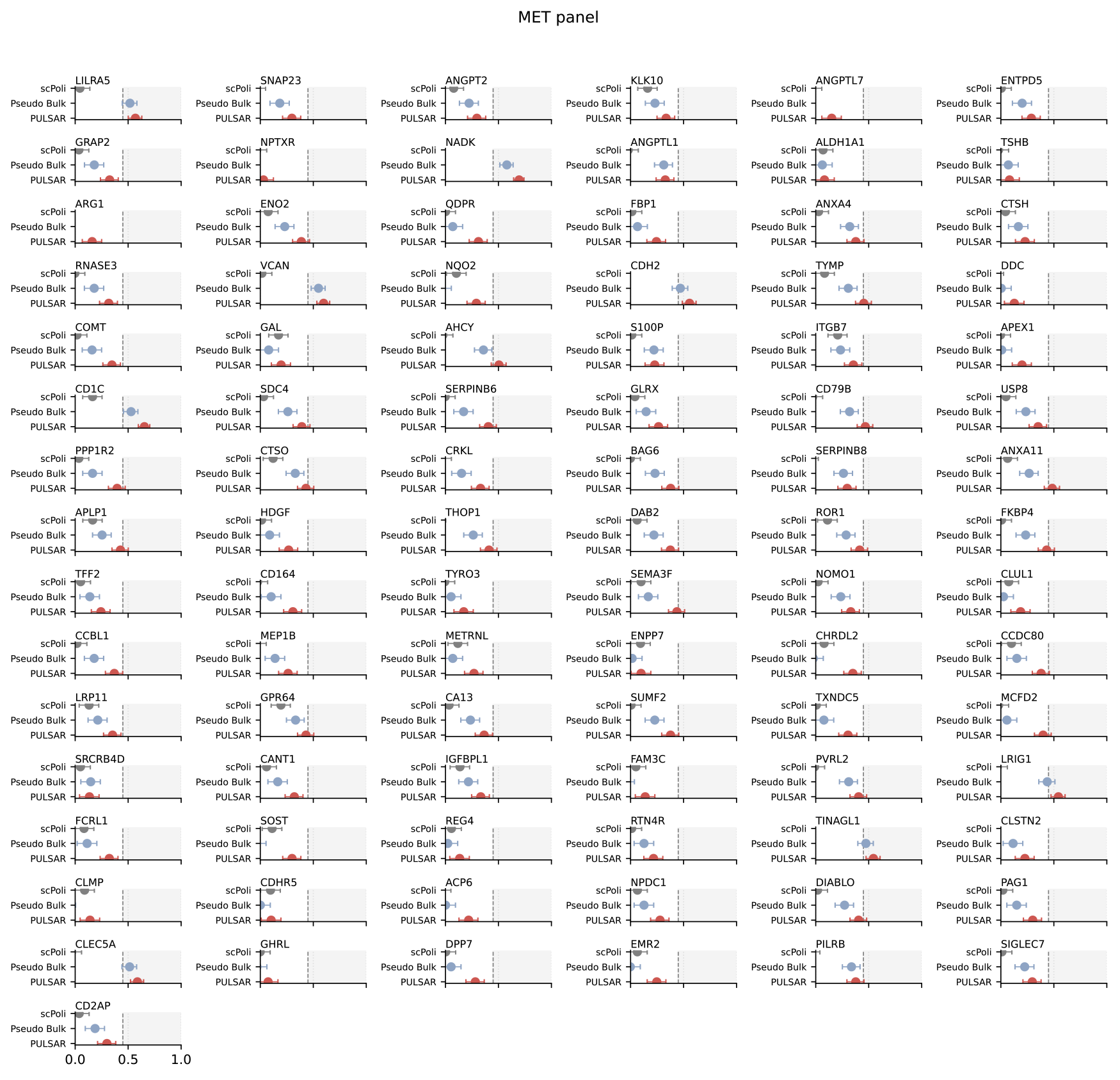
Protein prediction on the Olink MET panel. Per-analyte performance comparing scPoli, Pseudo-bulk, and PULSAR for predicting plasma protein abundance from baseline PBMC profiles. Points denote the mean Spearman correlation between predicted and measured levels across cross-validation folds; horizontal bars indicate 95% CI. The vertical reference line marks the performance threshold at a Spearman correlation of 0.45.

**Figure S9.**
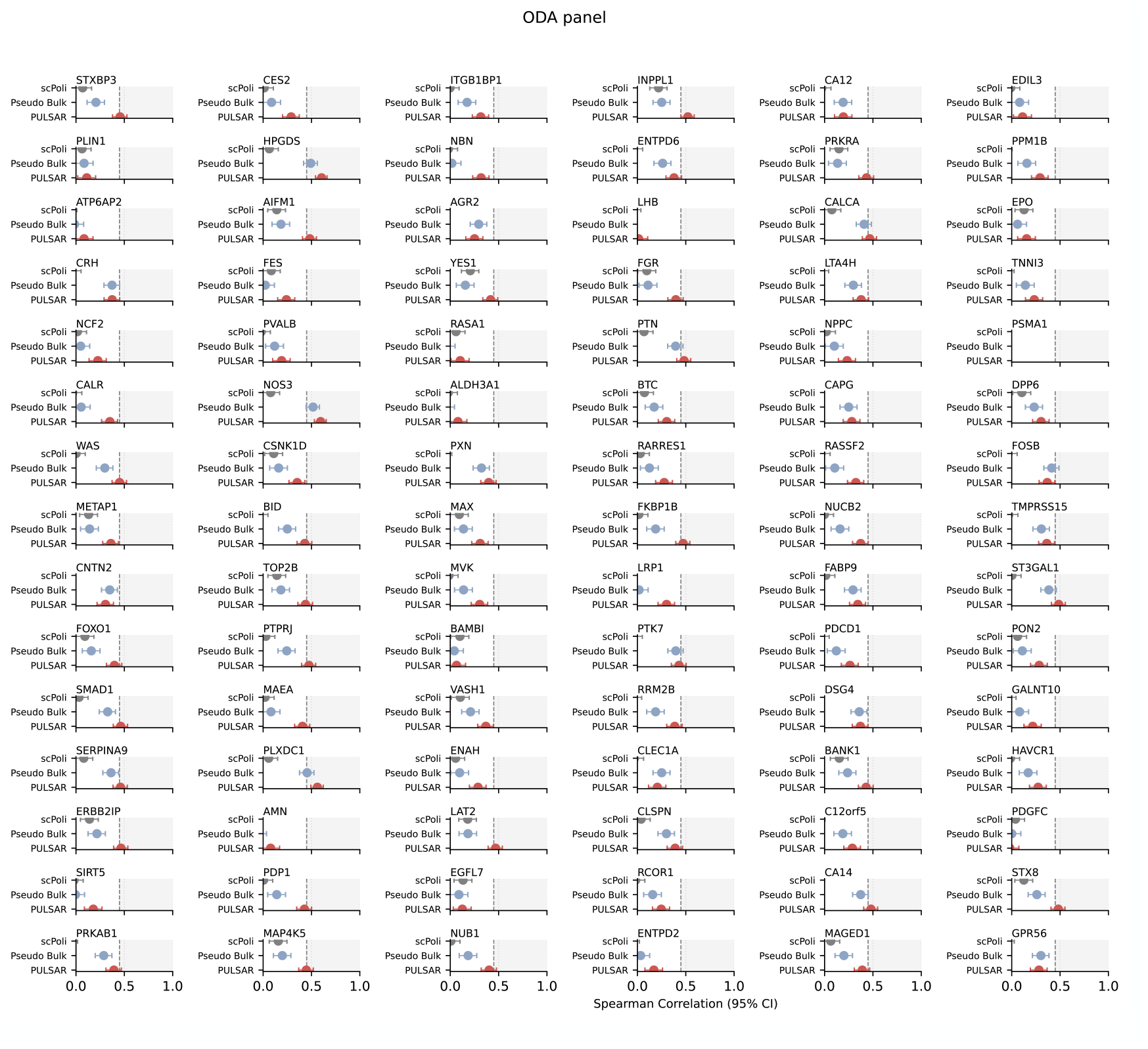
Protein prediction on the Olink ODA panel. Per-analyte performance comparing scPoli, Pseudo-bulk, and PULSAR for predicting plasma protein abundance from baseline PBMC profiles. Points denote the mean Spearman correlation between predicted and measured levels across cross-validation folds; horizontal bars indicate 95% CI. The vertical reference line marks the performance threshold at a Spearman correlation of 0.45.

**Figure S10.**
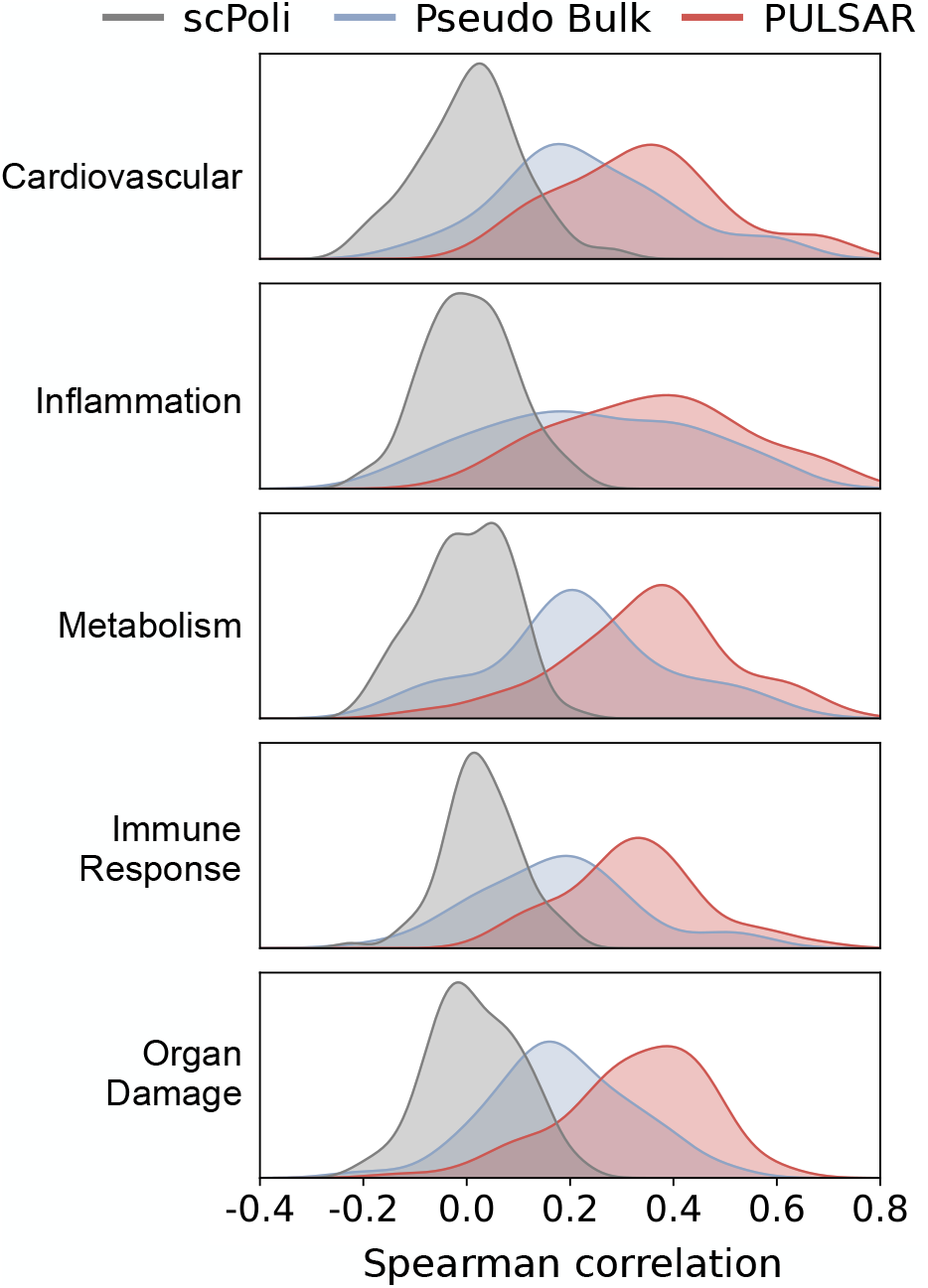
PULSAR improves protein prediction across all Olink panels. Kernel density estimates of per-analyte Spearman correlations between predicted and measured plasma protein levels for scPoli (grey), pseudobulk (blue), and PULSAR (red), stratified by Olink panel: Cardiovascular, Inflammation, Metabolism, Immune Response, and Organ Damage. Each curve summarizes the distribution of correlations for proteins within a panel; the consistent rightward shift of the PULSAR distributions indicates stronger predictive performance across all physiological domains.

**Figure S11.**
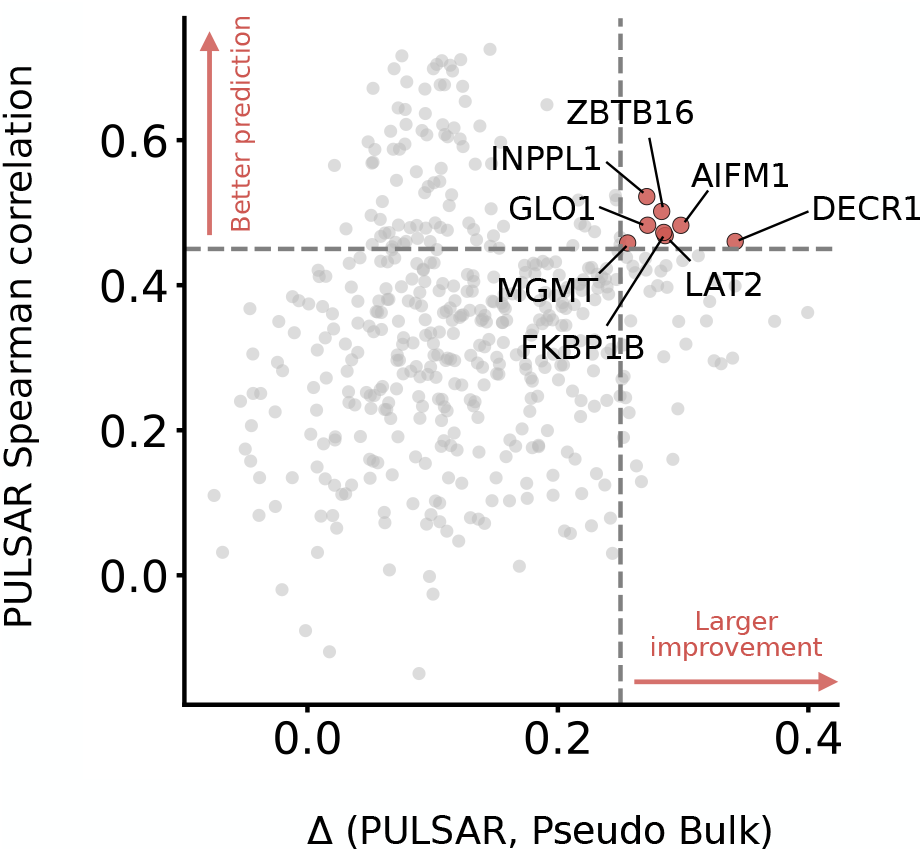
PULSAR most strongly improves prediction for proteins linked to discrete immune and metabolic programs. Each protein is plotted according to its PULSAR prediction performance (*y*-axis) and the improvement over pseudobulk (Δ(PULSAR, Pseudobulk), *x*-axis). We define thresholds on both axes to identify proteins where prediction accuracy improves significantly, transforming previously unpredictable targets into well-predicted ones. Proteins in the upper-left quadrant (PULSAR Spearman *r >* 0.45 and Δ(PULSAR, Pseudobulk) *<* 0.35) are well-predicted by both methods, indicating that bulk expression profiles suffice for their prediction. In contrast, proteins in the upper-right quadrant (highlighted in red dots, PULSAR Spearman *r* ≥ 0.45 and Δ(PULSAR, Pseudobulk) ≥ 0.35) show substantial improvement with PULSAR, achieving both high absolute performance and large gains over pseudobulk. Accordingly, markers linked to discrete immune subsets (e.g., ZBTB16, LAT2) and signaling adaptors (INPPL1, FKBP1B) benefit when the model captures shifts in rare populations and within–cell-type states. Likewise, mitochondrial and metabolic enzymes (AIFM1, DECR1, GLO1) and DNA repair machinery (MGMT) point to state-dependent release or turnover associated with oxidative stress, apoptosis, and proliferation. These patterns suggest that PULSAR’s advantage stems from preserving multicellular expression signatures that correlate with systemic protein levels.

**Figure S12.**
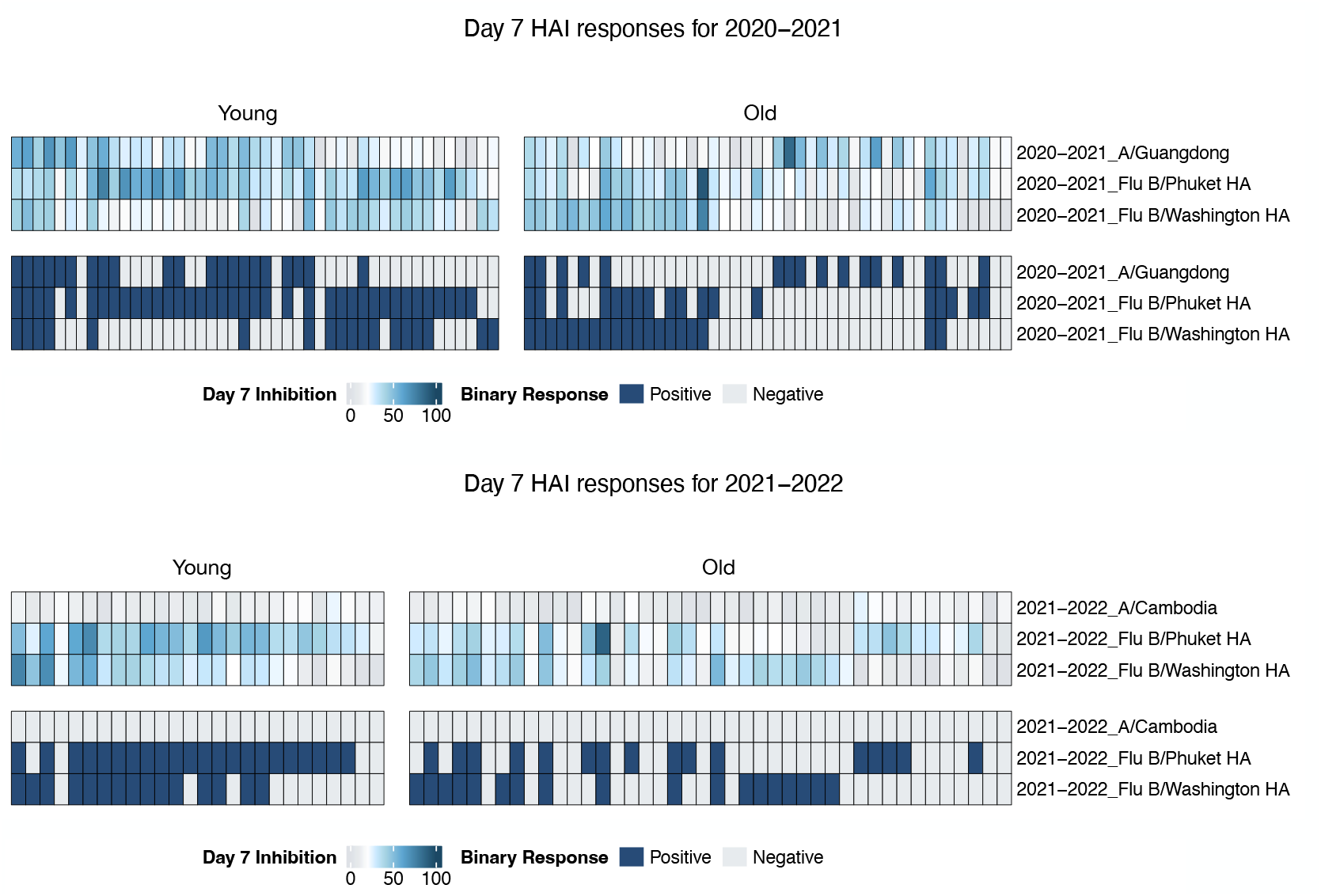
Day-7 hemagglutination inhibition (HAI) responses by age and season. Heat maps show individual day-7 HAI percentage inhibition for three vaccine strains in two influenza seasons (2020–2021 and 2021–2022), stratified into Young and Older cohorts. For each strain, the upper band displays inhibition titres (0–100; light to dark blue), and the lower band shows the corresponding binary responder status (dark blue = positive, grey = negative).

**Figure S13.**
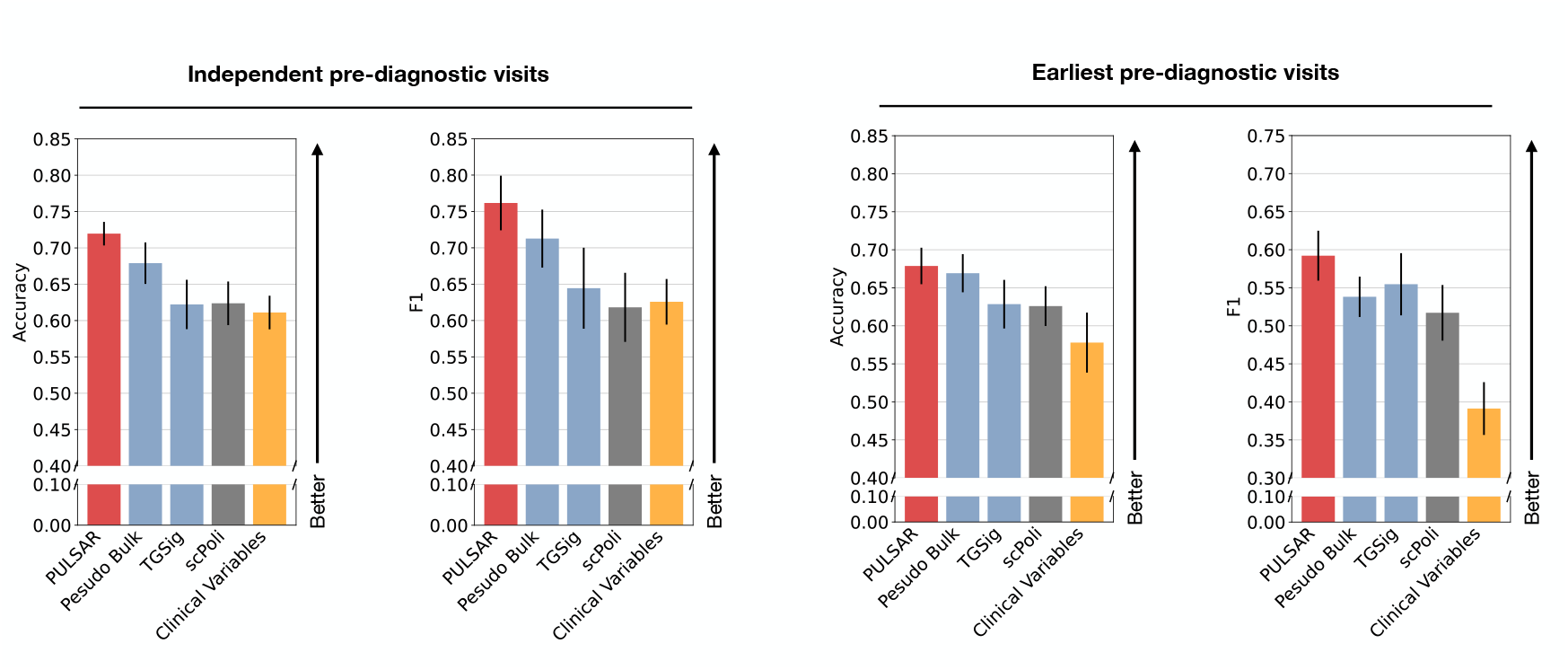
PULSAR improves prediction of rheumatoid arthritis conversion from baseline PBMC profiles. Bar plots show mean Accuracy (left) and F1 score (right) for predicting progression from ACPA^+^ at-risk status to clinical rheumatoid arthritis, comparing PULSAR, pseudobulk, TGSig, scPoli, and a clinical-variable model. Performance is shown for (left pair) all independent pre-diagnostic visits treated as separate samples and (right pair) the earliest available pre-diagnostic visit per donor. Error bars indicate s.e.m. across repeated stratified cross-validation folds. PULSAR consistently achieves the highest accuracy and F1 in both settings, outperforming transcriptomic baselines and clinical variables alone.

**Figure S14.**
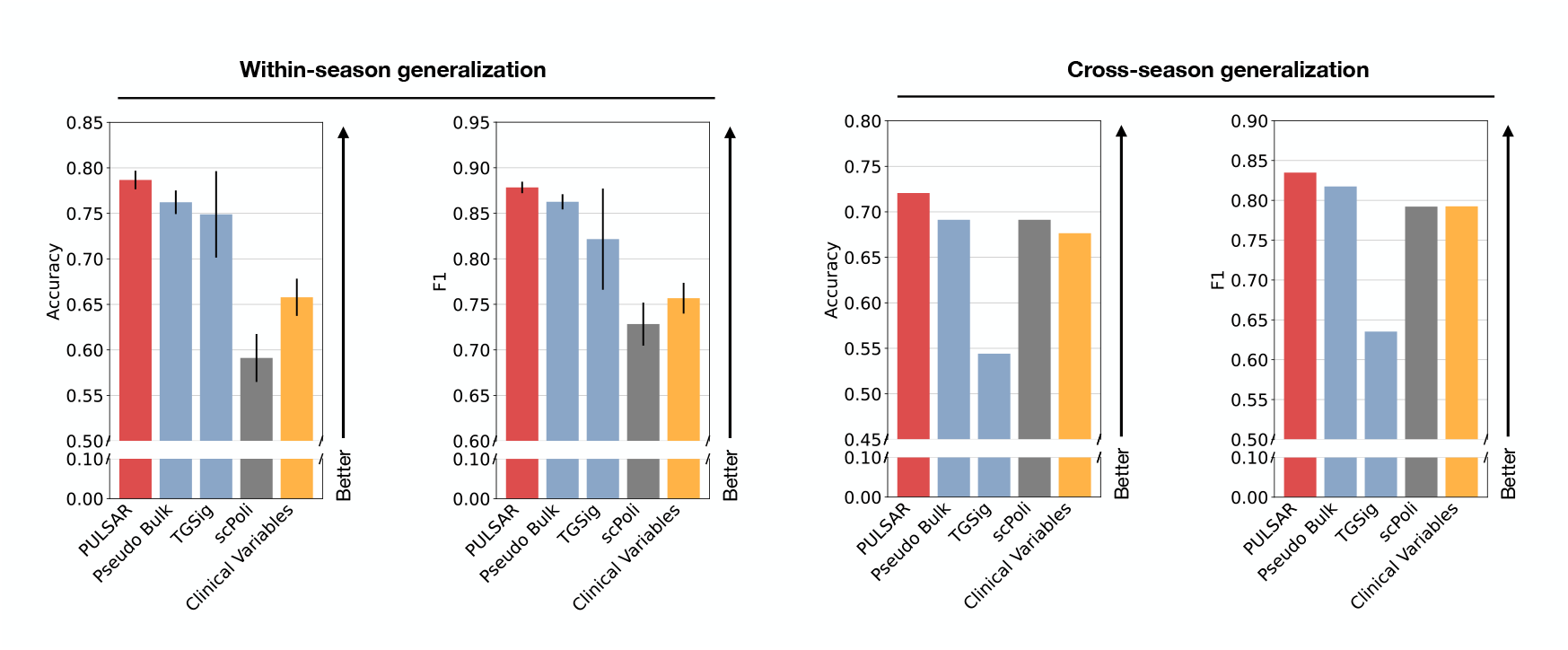
PULSAR predicts influenza vaccine responsiveness within and across seasons from baseline PBMC profiles. Bar plots show mean Accuracy (left) and F1 score (right) for classifying high versus low responders to seasonal influenza vaccination using day-0 PBMC scRNA-seq. Methods compared are PULSAR, pseudobulk, TGSig, scPoli, and a clinical-variable model. Performance is shown for within-season generalization (training and testing on the same flu season; left pair) and cross-season generalization (training on one season and testing on the next; right pair). Error bars indicate s.e.m. across repeated stratified cross-validation. PULSAR consistently achieves the highest accuracy and F1 in both settings, outperforming transcriptomic baselines and models using clinical variables alone.

**Figure S15.**
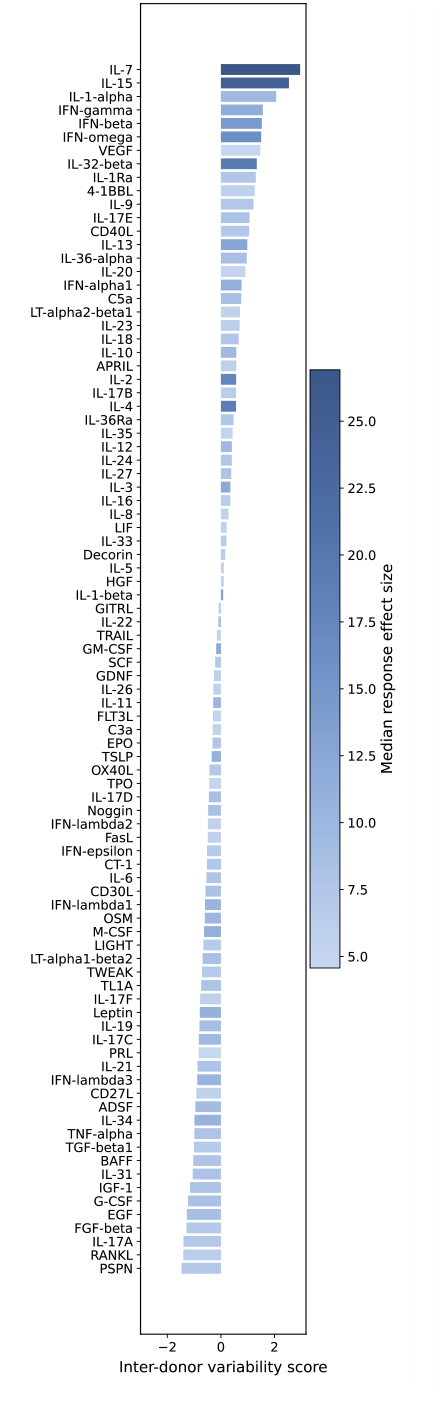
Cytokine-specific individualization score of donor responses. Ranked bar plot of the individualized response score for each of 90 cytokines. Colored by the median effect size in PULSAR donor embedding space.

**Figure S16.**
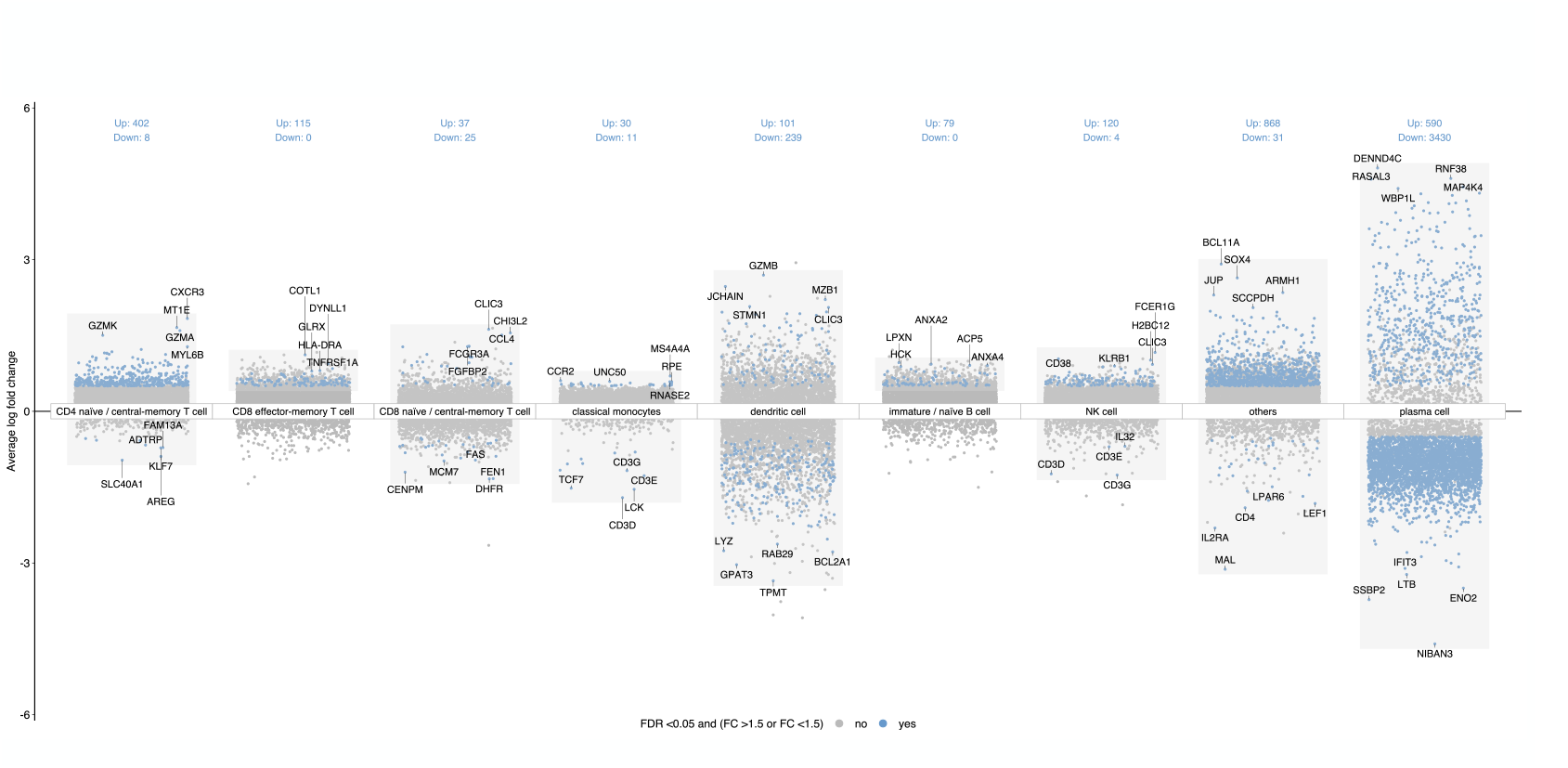
Cell type–resolved differential expression across condition and control. Scatter panels show per-gene average log fold change (y-axis) between the condition and matched controls, stratified by major PBMC cell types as annotated by scRNA-seq. Each point is a gene; blue points denote significantly differentially expressed genes (FDR *<* 0.05 and |Fold change| *>* 1.5), and a subset of sentinel genes is labeled for orientation. Numbers above each panel indicate counts of up-regulated (Up) and down-regulated (Down) genes in that cell type.

**Table S1.**
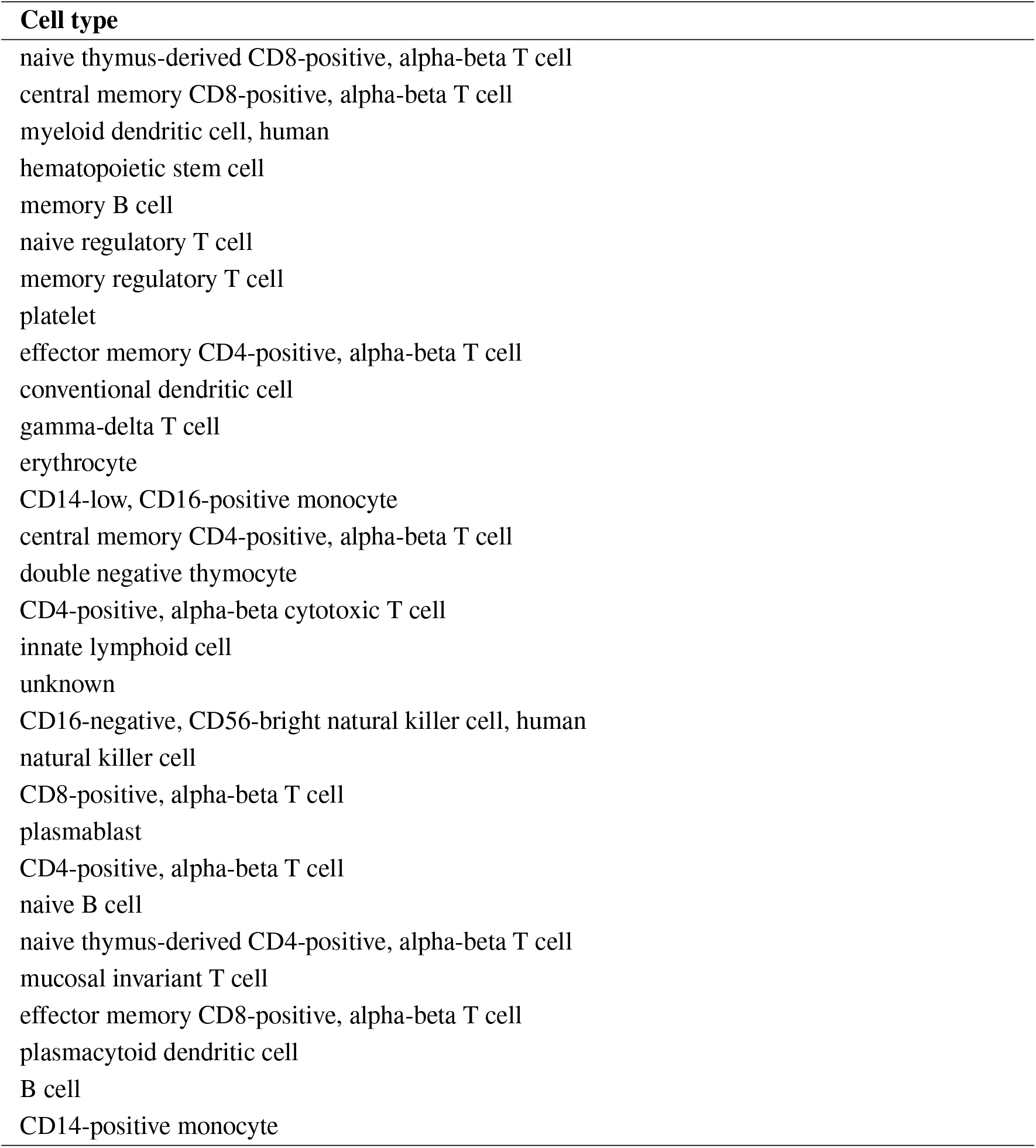
Cell type list for constructing the cell-type-proportion baseline.

