## Supplementary Information for "PULSAR: a Foundation Model for Multi-scale and Multicellular Biology"


This PDF file includes:

- Supplementary Figure 1-16
- Supplementary Table 1

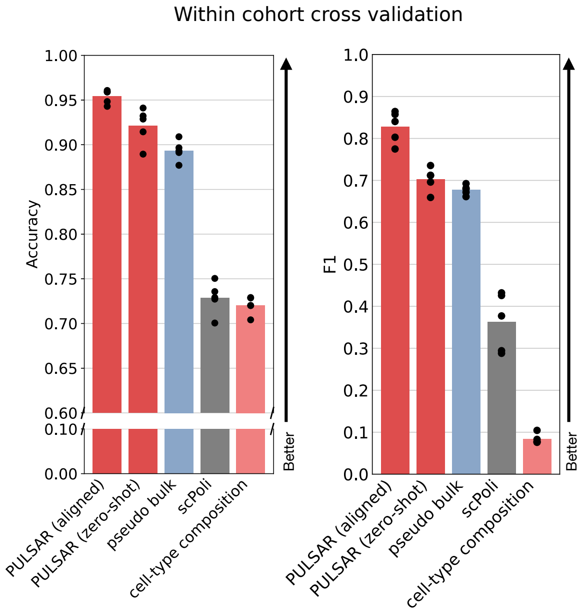


**Figure S1: Performance benchmark for within-cohort cross-validation of disease classification** Accuracy (left) and F1 (right). Points denote the mean over 5-fold runs.

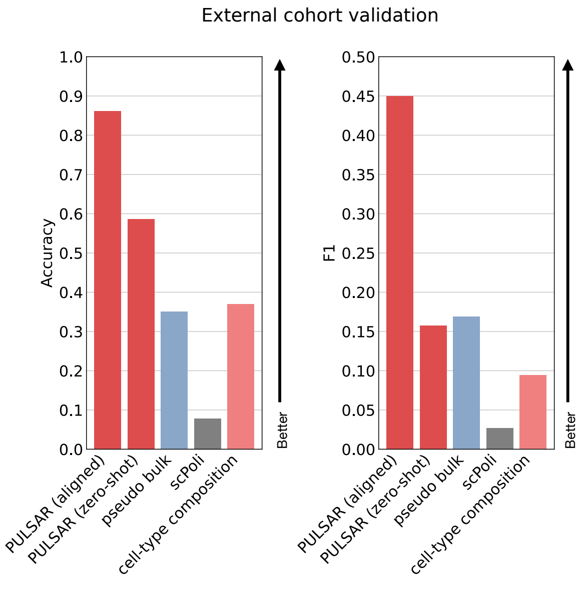


**Figure S2: Performance benchmark for external cohort cross-validation of disease classification** Accuracy (left) and F1 (right). Points denote the mean over 5-fold runs.

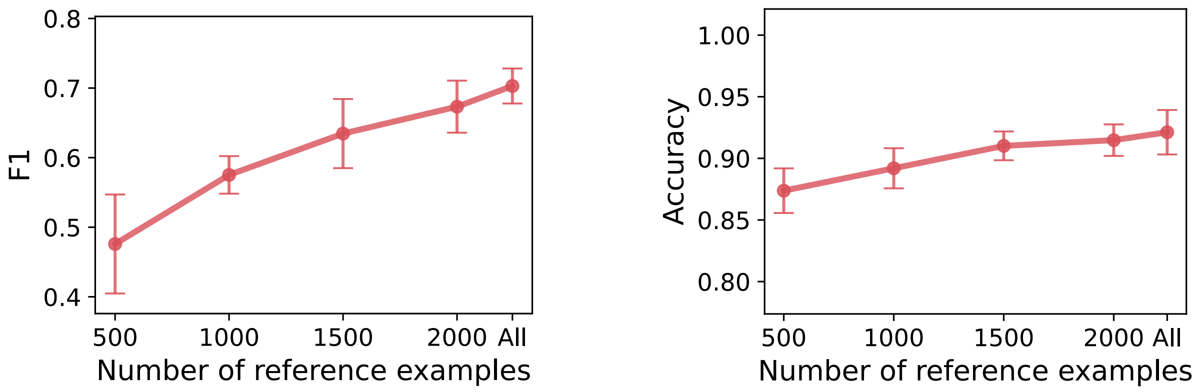


**Figure S3: Performance scales with the size of the reference vector database.** Macro F1 (left) and Accuracy (right) as a function of the number of reference examples retrieved from the vector database in the reference split. All is 2243 samples. Points denote the mean over 5-fold runs, error bars indicate s.e.m. We observed a scaling effect as the size of the reference vector database increased. Expanding from 500 to 1,500 reference examples yielded the largest gains (Macro F1 from 0.588 to 0.669; Accuracy 0.875 to 0.891), with steady improvements up to the full reference corpus (Macro F1 0.712; Accuracy 0.889). The trends indicate that expanding the reference corpus improves search accuracy.

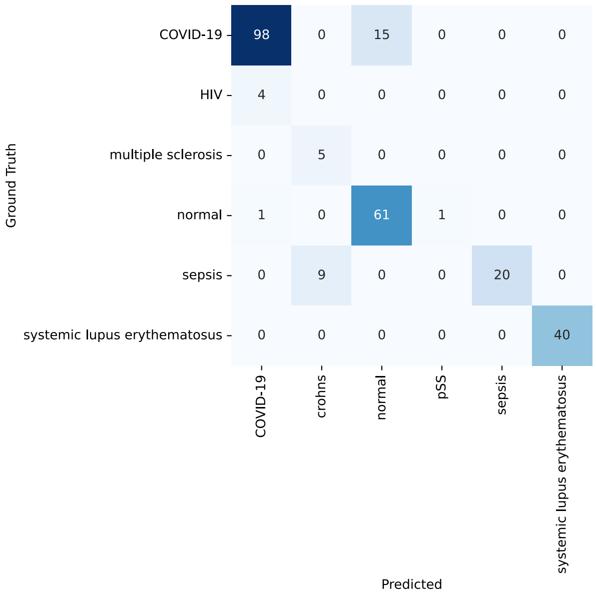


**Figure S4: Confusion matrix on the external cohort.** Confusion matrix summarizing predictions across an external cohort at the disease level.

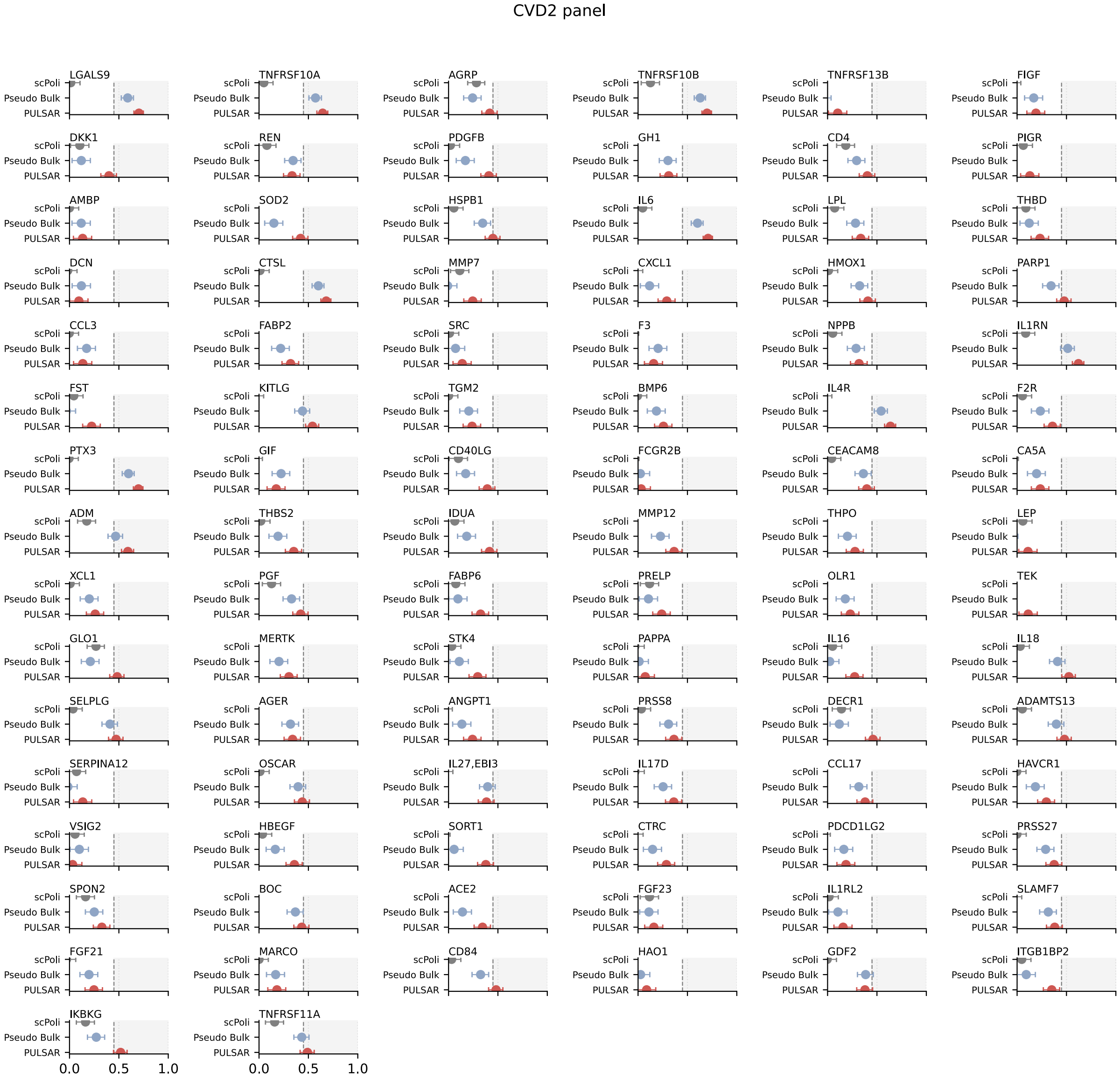


**Figure S5: Protein prediction on the Olink CVD2 panel.** Per-analyte performance comparing scPoli, Pseudo-bulk, and PULSAR for predicting plasma protein abundance from baseline PBMC profiles. Points denote the mean Spearman correlation between predicted and measured levels across cross-validation folds; horizontal bars indicate 95% CI. The vertical reference line marks the performance threshold at a Spearman correlation of 0.45.

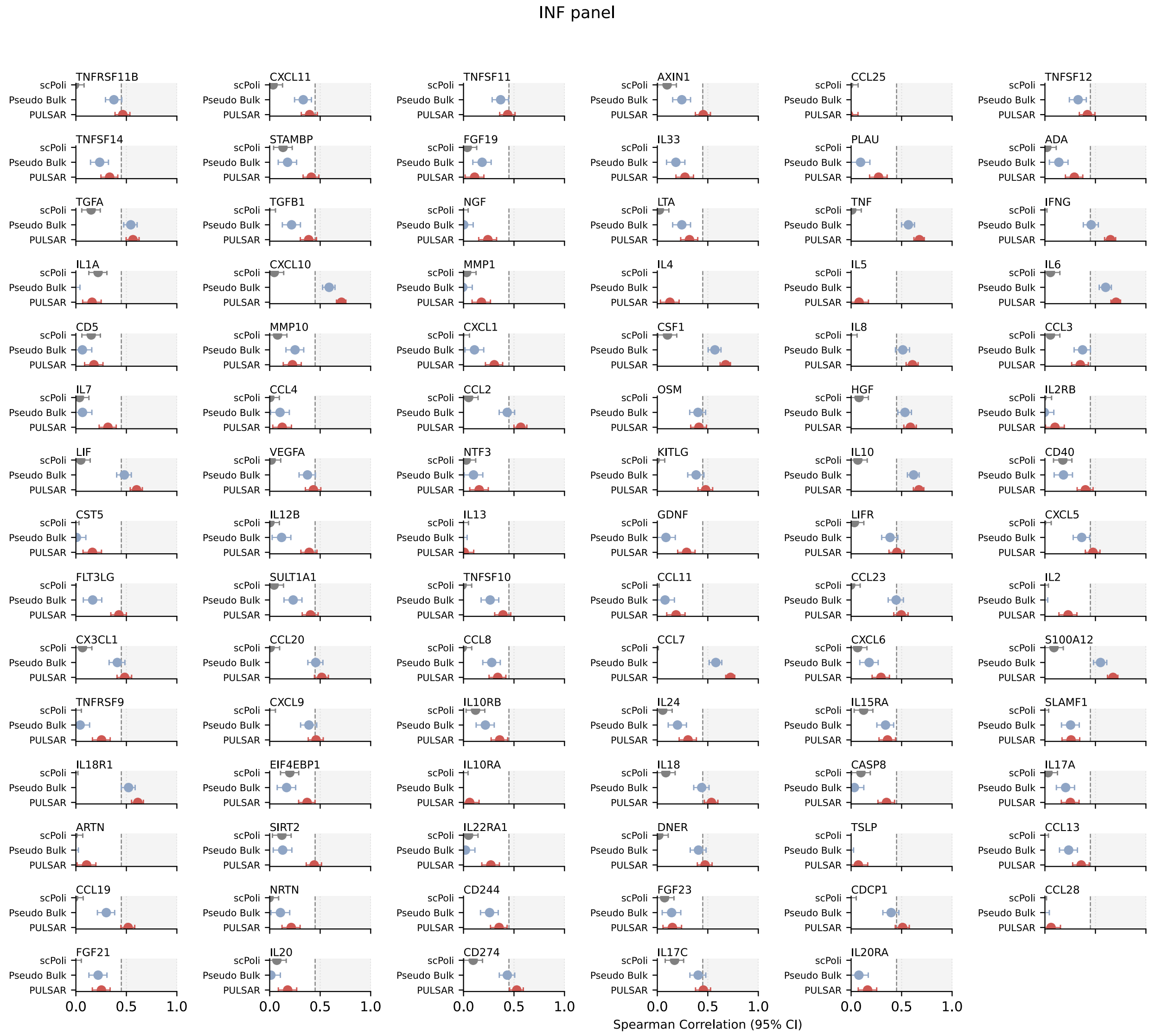


**Figure S6: Protein prediction on the Olink INF panel.** Per-analyte performance comparing scPoli, Pseudo-bulk, and PULSAR for predicting plasma protein abundance from baseline PBMC profiles. Points denote the mean Spearman correlation between predicted and measured levels across cross-validation folds; horizontal bars indicate 95% CI. The vertical reference line marks the performance threshold at a Spearman correlation of 0.45.

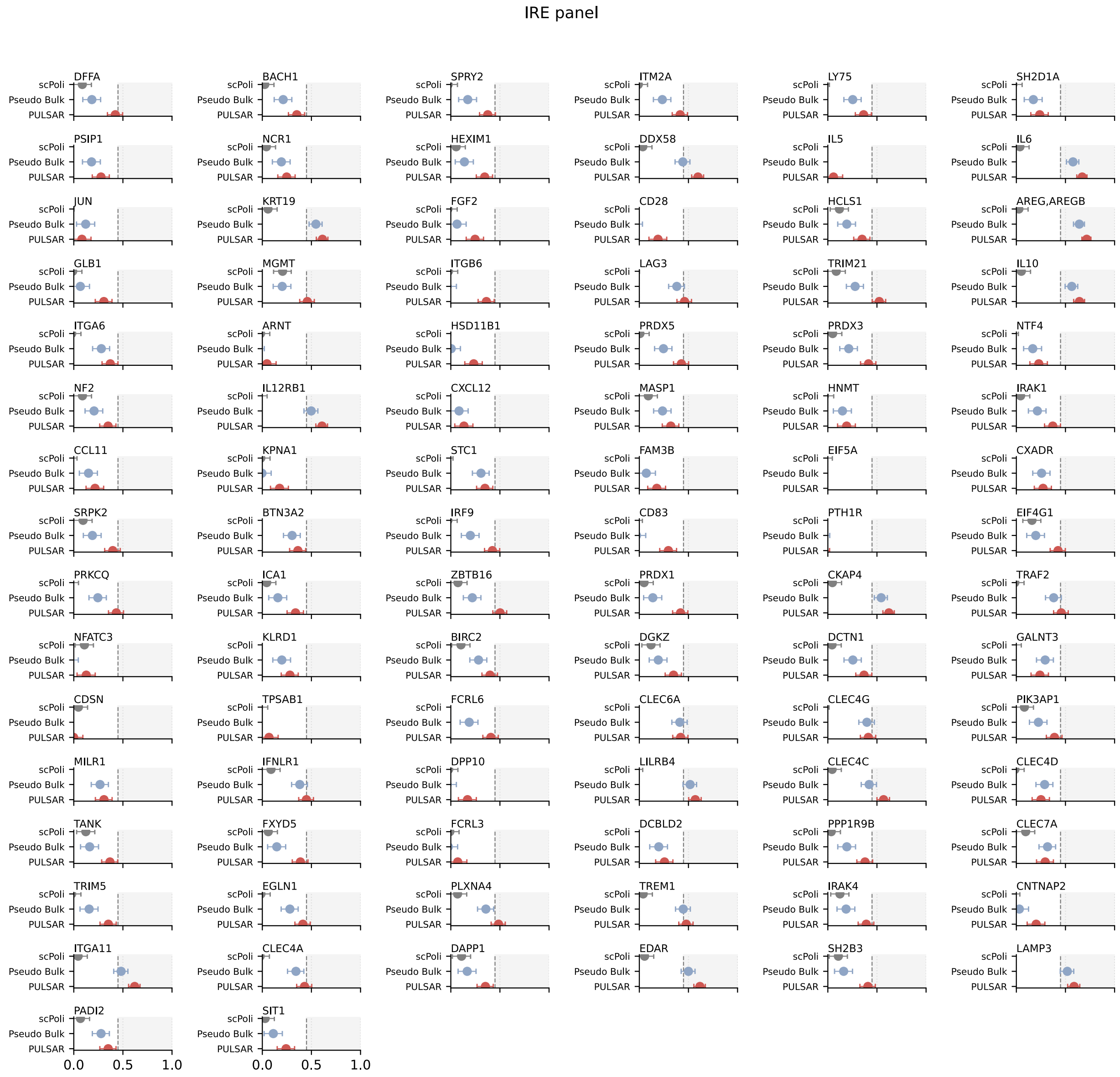


**Figure S7: Protein prediction on the Olink IRE panel.** Per-analyte performance comparing scPoli, Pseudo-bulk, and PULSAR for predicting plasma protein abundance from baseline PBMC profiles. Points denote the mean Spearman correlation between predicted and measured levels across cross-validation folds; horizontal bars indicate 95% CI. The vertical reference line marks the performance threshold at a Spearman correlation of 0.45.

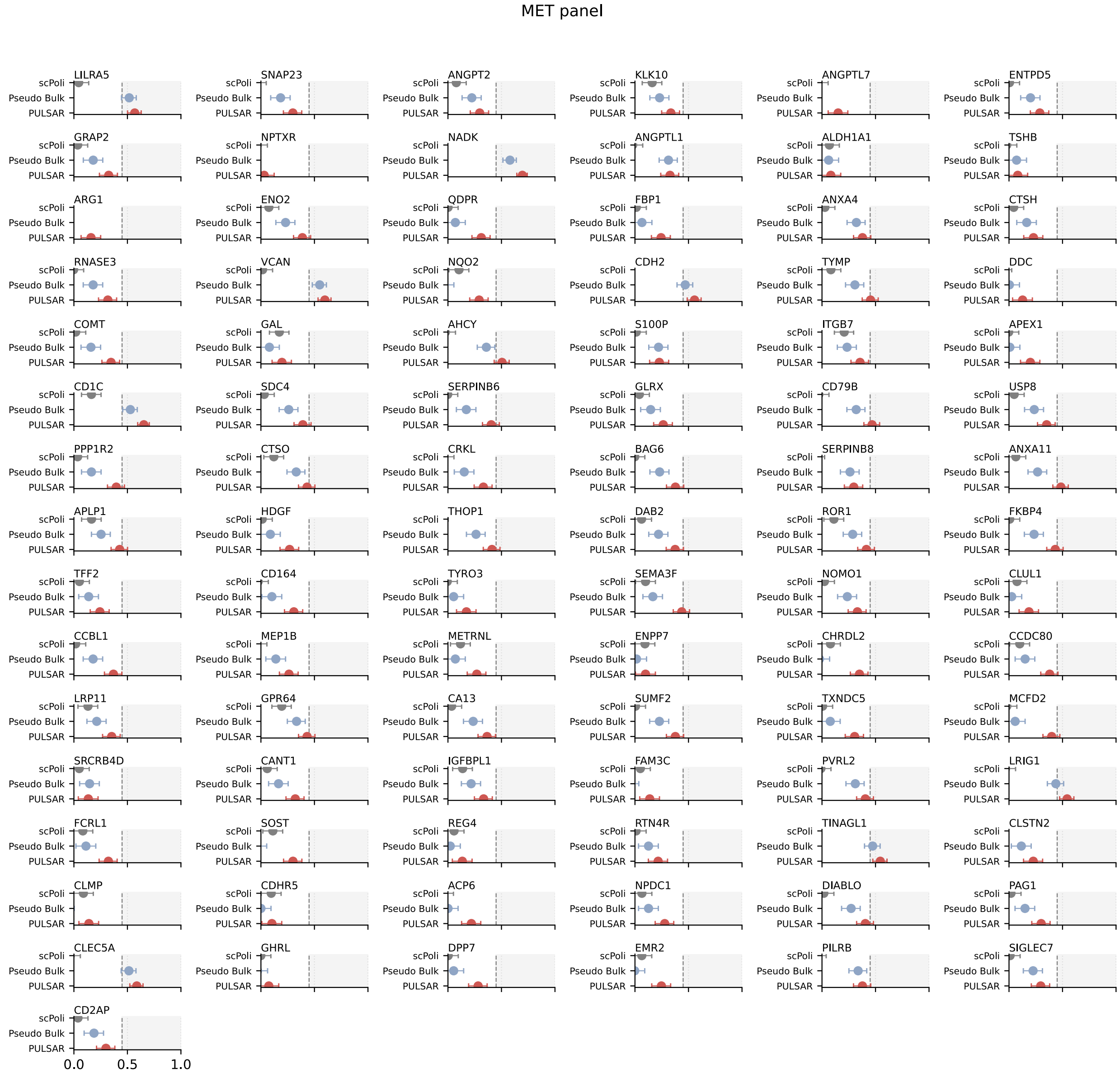


**Figure S8: Protein prediction on the Olink MET panel.** Per-analyte performance comparing scPoli, Pseudo-bulk, and PULSAR for predicting plasma protein abundance from baseline PBMC profiles. Points denote the mean Spearman correlation between predicted and measured levels across cross-validation folds; horizontal bars indicate 95% CI. The vertical reference line marks the performance threshold at a Spearman correlation of 0.45.

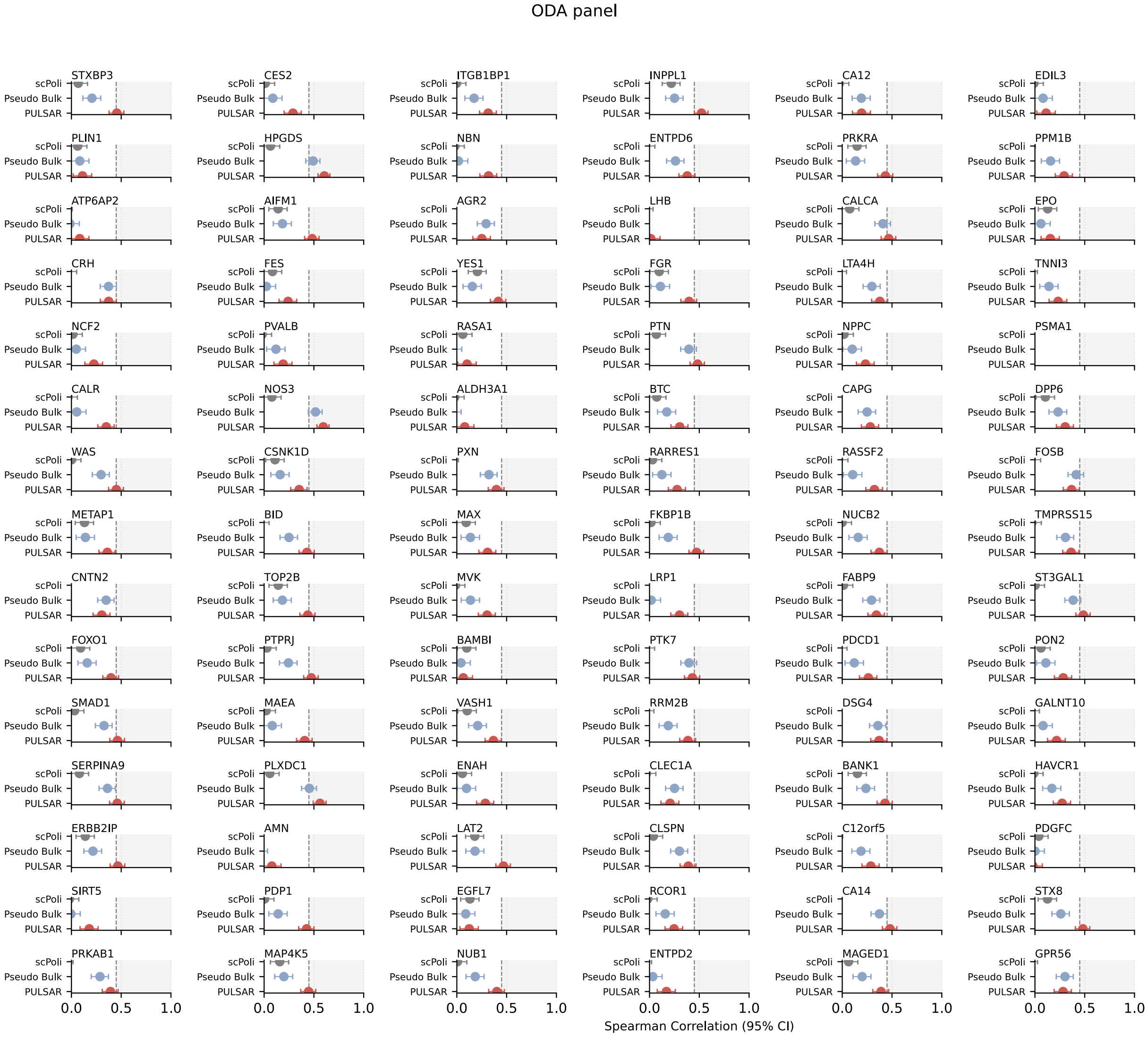


**Figure S9: Protein prediction on the Olink ODA panel.** Per-analyte performance comparing scPoli, Pseudo-bulk, and PULSAR for predicting plasma protein abundance from baseline PBMC profiles. Points denote the mean Spearman correlation between predicted and measured levels across cross-validation folds; horizontal bars indicate 95% CI. The vertical reference line marks the performance threshold at a Spearman correlation of 0.45.

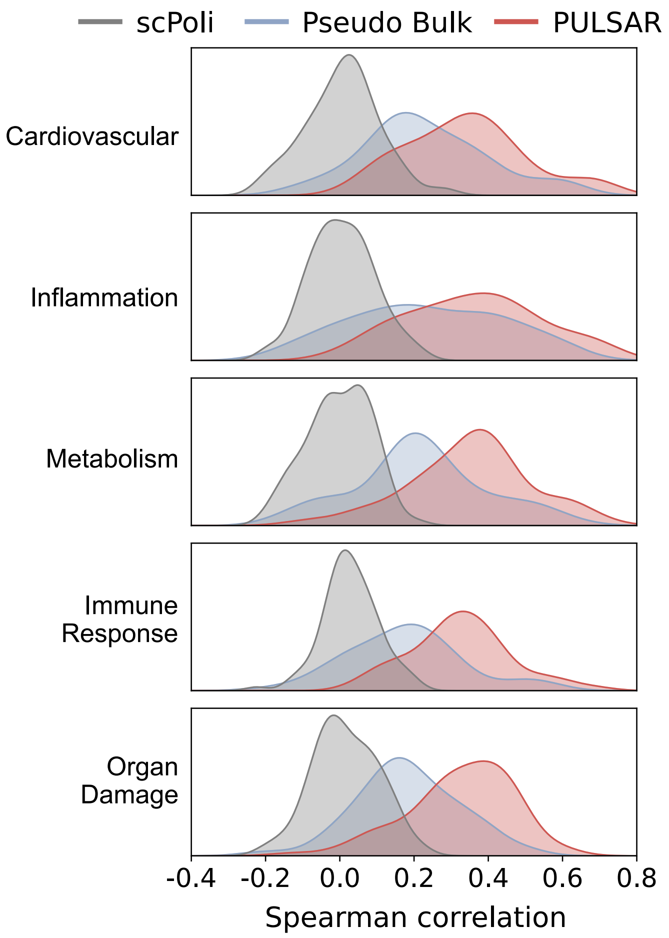


**Figure S10: PULSAR improves protein prediction across all Olink panels.** Kernel density estimates of per-analyte Spearman correlations between predicted and measured plasma protein levels for scPoli (grey), pseudobulk (blue), and PULSAR (red), stratified by Olink panel: Cardiovascular, Inflammation, Metabolism, Immune Response, and Organ Damage. Each curve summarizes the distribution of correlations for proteins within a panel; the consistent rightward shift of the PULSAR distributions indicates stronger predictive performance across all physiological domains.

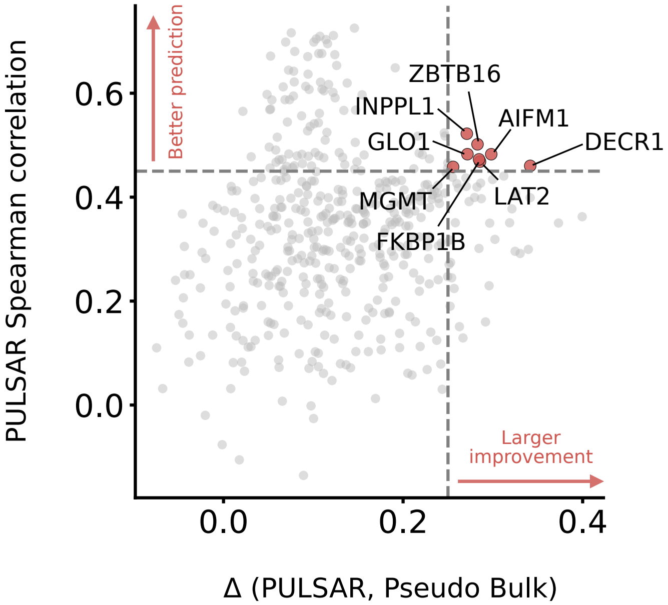


**Figure S11: PULSAR most strongly improves prediction for proteins linked to discrete immune and metabolic programs.** Each protein is plotted according to its PULSAR prediction performance ($y$-axis) and the improvement over pseudobulk (Δ(PULSAR, Pseudobulk), $x$-axis). We define thresholds on both axes to identify proteins where prediction accuracy improves significantly, transforming previously unpredictable targets into well-predicted ones. Proteins in the upper-left quadrant (PULSAR Spearman $r > 0.45$ and Δ(PULSAR, Pseudobulk) $< 0.35$) are well-predicted by both methods, indicating that bulk expression profiles suffice for their prediction. In contrast, proteins in the upper-right quadrant (highlighted in red dots, PULSAR Spearman $r \geq 0.45$ and Δ(PULSAR, Pseudobulk) $\geq$ 0.35) show substantial improvement with PULSAR, achieving both high absolute performance and large gains over pseudobulk. Accordingly, markers linked to discrete immune subsets (e.g., ZBTB16, LAT2) and signaling adaptors (INPPL1, FKBP1B) benefit when the model captures shifts in rare populations and within–cell-type states. Likewise, mitochondrial and metabolic enzymes (AIFM1, DECR1, GLO1) and DNA repair machinery (MGMT) point to state-dependent release or turnover associated with oxidative stress, apoptosis, and proliferation. These patterns suggest that PULSAR's advantage stems from preserving multicellular expression signatures that correlate with systemic protein levels.

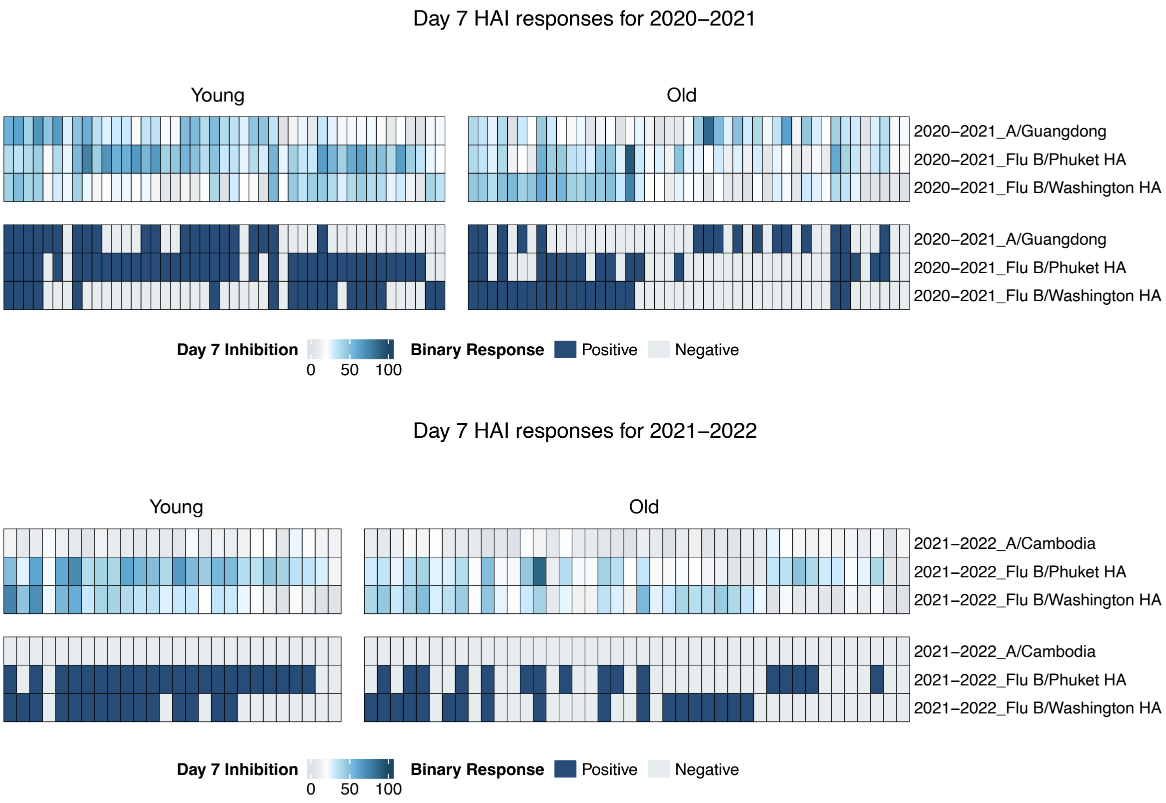


**Figure S12: Day-7 hemagglutination inhibition (HAI) responses by age and season.** Heat maps show individual day-7 HAI percentage inhibition for three vaccine strains in two influenza seasons (2020–2021 and 2021–2022), stratified into Young and Older cohorts. For each strain, the upper band displays inhibition titres (0–100; light to dark blue), and the lower band shows the corresponding binary responder status (dark blue = positive, grey = negative).

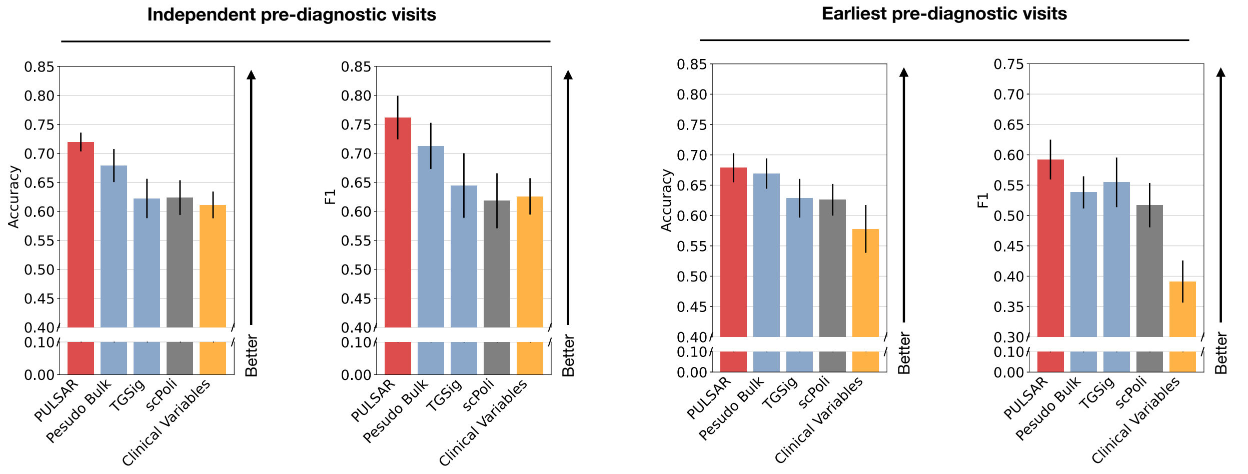


**Figure S13: PULSAR improves prediction of rheumatoid arthritis conversion from baseline PBMC profiles.** Bar plots show mean Accuracy (left) and F1 score (right) for predicting progression from $ACPA^+$ at-risk status to clinical rheumatoid arthritis, comparing PULSAR, pseudobulk, TGSig, scPoli, and a clinical-variable model. Performance is shown for (left pair) all independent pre-diagnostic visits treated as separate samples and (right pair) the earliest available pre-diagnostic visit per donor. Error bars indicate s.e.m. across repeated stratified cross-validation folds. PULSAR consistently achieves the highest accuracy and F1 in both settings, outperforming transcriptomic baselines and clinical variables alone.

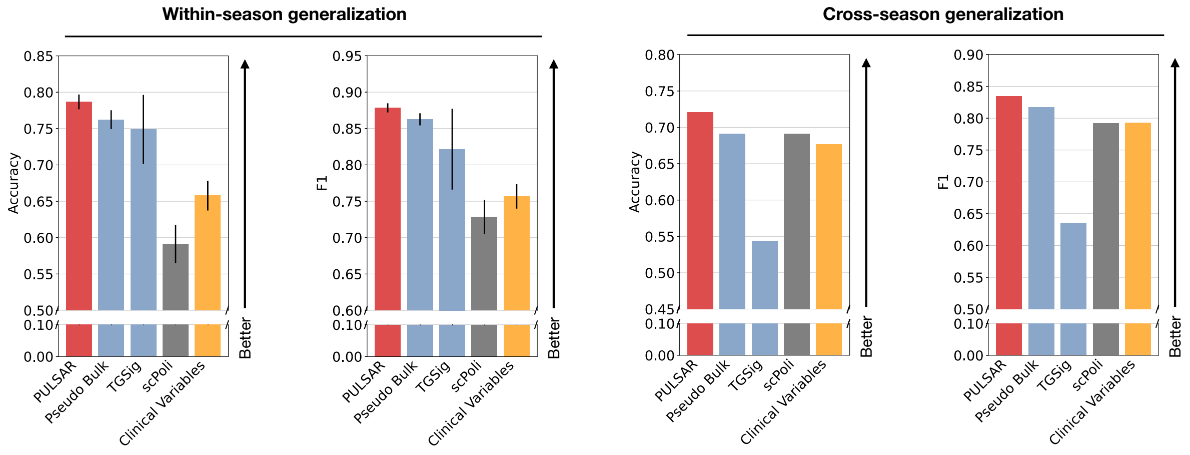


**Figure S14: PULSAR predicts influenza vaccine responsiveness within and across seasons from baseline PBMC profiles.** Bar plots show mean Accuracy (left) and F1 score (right) for classifying high versus low responders to seasonal influenza vaccination using day-0 PBMC scRNA-seq. Methods compared are PULSAR, pseudobulk, TGSig, scPoli, and a clinical-variable model. Performance is shown for within-season generalization (training and testing on the same flu season; left pair) and cross-season generalization (training on one season and testing on the next; right pair). Error bars indicate s.e.m. across repeated stratified cross-validation. PULSAR consistently achieves the highest accuracy and F1 in both settings, outperforming transcriptomic baselines and models using clinical variables alone.

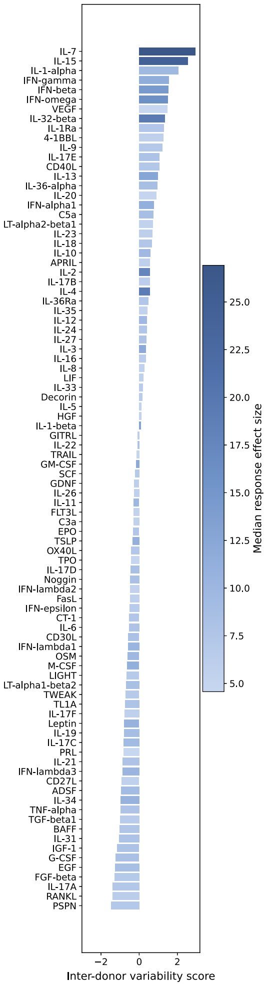


**Figure S15: Cytokine-specific individualization score of donor responses.** Ranked bar plot of the individualized response score for each of 90 cytokines. Colored by the median effect size in PULSAR donor embedding space.

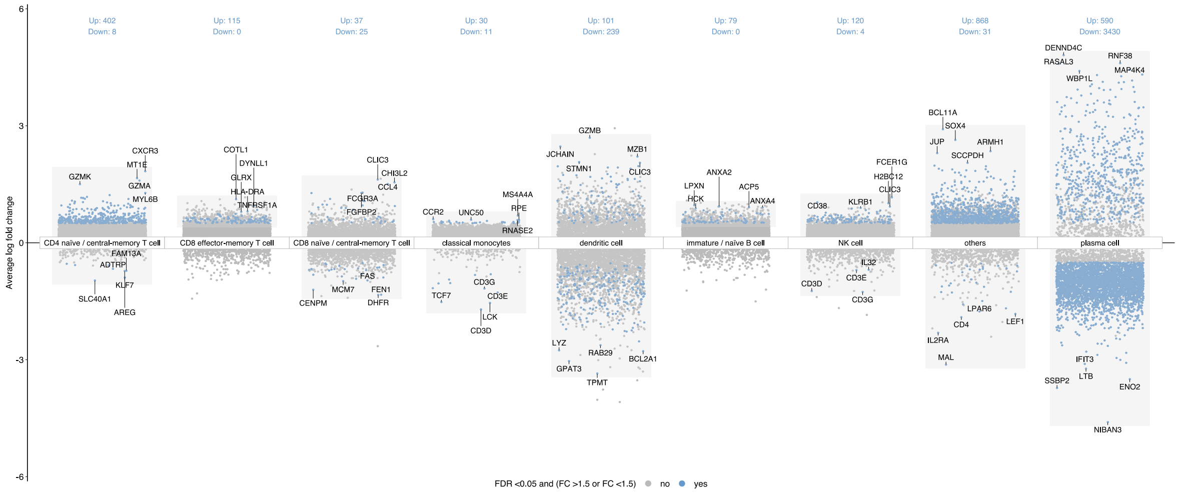


**Figure S16: Cell type–resolved differential expression across condition and control.** Scatter panels show per-gene average log fold change (y-axis) between the condition and matched controls, stratified by major PBMC cell types as annotated by scRNA-seq. Each point is a gene; blue points denote significantly differentially expressed genes (FDR $< 0.05$ and $|\text{Fold change}| > 1.5$), and a subset of sentinel genes is labeled for orientation. Numbers above each panel indicate counts of up-regulated (Up) and down-regulated (Down) genes in that cell type.

| **Cell type** |
|---|
| naive thymus-derived CD8-positive, alpha-beta T cell |
| central memory CD8-positive, alpha-beta T cell |
| myeloid dendritic cell, human |
| hematopoietic stem cell |
| memory B cell |
| naive regulatory T cell |
| memory regulatory T cell |
| platelet |
| effector memory CD4-positive, alpha-beta T cell |
| conventional dendritic cell |
| gamma-delta T cell |
| erythrocyte |
| CD14-low, CD16-positive monocyte |
| central memory CD4-positive, alpha-beta T cell |
| double negative thymocyte |
| CD4-positive, alpha-beta cytotoxic T cell |
| innate lymphoid cell |
| unknown |
| CD16-negative, CD56-bright natural killer cell, human |
| natural killer cell |
| CD8-positive, alpha-beta T cell |
| plasmablast |
| CD4-positive, alpha-beta T cell |
| naive B cell |
| naive thymus-derived CD4-positive, alpha-beta T cell |
| mucosal invariant T cell |
| effector memory CD8-positive, alpha-beta T cell |
| plasmacytoid dendritic cell |
| B cell |
| CD14-positive monocyte |

**Table S1:** Cell type list for constructing the cell-type-proportion baseline.
